# Sex-biased islet β cell dysfunction is caused by the MODY MAFA S64F variant by inducing premature aging and senescence in males

**DOI:** 10.1101/2020.06.29.177527

**Authors:** Emily M. Walker, Jeeyeon Cha, Xin Tong, Min Guo, Jin-Hua Liu, Sophia Yu, Donato Iacovazzo, Franck Mauvais-Jarvis, Sarah E. Flanagan, Márta Korbonits, John Stafford, David Jacobson, Roland Stein

## Abstract

A heterozygous missense mutation producing a variant of the islet β-cell-enriched MAFA transcription factor (Ser(S)64Phe(F) MAFA) was identified in humans who developed adult-onset, β-cell dysfunction (diabetes or insulinomatosis), with men more prone to diabetes. This mutation engenders increased stability to the normally unstable MAFA protein. To obtain insight into how this variant impacts β cell function, we developed a mouse model expressing S64F MafA and found sex-dependent phenotypes, with heterozygous mutant males displaying impaired glucose tolerance while females were slightly hypoglycemic with improved blood glucose clearance. Only heterozygous males showed transiently higher MafA protein levels preceding the onset of glucose intolerance and sex-dependent, differential expression of genes involved in calcium signaling, DNA damage, aging, and senescence. Functional changes in islet calcium handling and signs of islet aging and senescence processes were uniquely observed in male animals. In addition, S64F MAFA expression in human, male EndoC-βH2 β cells accelerated cellular senescence and increased production of senescence-associated secretory proteins compared to cells expressing wild-type MAFA. Together, these results implicate a conserved mechanism of accelerated islet aging and senescence in promoting diabetes in S64F MAFA carriers in a sex-dependent manner.

## Introduction

The large V-<u>M</u>af <u>A</u>vian Musculoaponeurotic <u>F</u>ibrosarcoma transcription factor Maf<u>A</u> is a pancreatic β-cell-enriched protein essential for activating rodent transcriptional programs to promote cell maturation and glucose-stimulated insulin secretion (GSIS) (Artner et al., 2008; Artner et al., 2010; Hang and Stein, 2011; Hang et al., 2014; Zhang et al., 2005). The presence of MafA in only insulin^+^ β cells during islet cell development and postnatally distinguishes it from other important islet cell-enriched regulators (Golson and Kaestner, 2017; Hang and Stein, 2011; Pan and Wright, 2011; Shih et al., 2013). For example, transcription factors such as Pdx1 and Nkx6.1, which are expressed earlier and more broadly, also have a profound impact on pancreas organogenesis, islet β cell function, and whole body glucose homeostasis in mice (Madsen et al., 1997; Pan and Wright, 2011). Pdx1 is present in both the exocrine and endocrine pancreas during development and then principally in islet β cells postnatally (Pan and Wright, 2011), while Nkx6.1 is found in endocrine progenitors and is later restricted to β cells (Golson and Kaestner, 2017; Hang and Stein, 2011; Pan and Wright, 2011; Shih et al., 2013). In contrast, mice lacking MafA (i.e. *MafA^−/−^* (Zhang et al., 2005), *MafA*^*Δpanc*^ (Artner et al., 2010; Hang et al., 2014), *MafA*^*Δ*β^ (Cyphert et al., 2019; Luan et al., 2019)) have normal pancreas formation and have a relatively subtle influence on postnatal physiology, primarily compromising GSIS in a sex-independent manner. However, MafA has been proposed to play a distinct late role in postnatal islet β cell maturation, in part supported by observations demonstrating that inducing MafA expression levels in normally non-glucose responsive and MafA^Low^ neonatal rat islets increases GSIS (Aguayo-Mazzucato et al., 2011), while compromising levels reduces mouse (Artner et al., 2010; Cyphert et al., 2019; Hang et al., 2014; Luan et al., 2019; Zhang et al., 2005) and human (Guo et al., 2013) β cell activity. A key regulatory role is also implied by the ability of many islet-enriched transcription factors to coordinately stimulate *MafA* gene transcription, including Pdx1 and Nkx6.1 (Raum et al., 2006; Raum et al., 2010).

Expression of the other large Maf family member produced in the islet, MafB, differs from MafA in being produced in both murine α- and β-cells developmentally (Cyphert et al., 2019; Hang and Stein, 2011), then postnatally only in α cells and a subpopulation of β cells during pregnancy (Banerjee et al., 2016; Cyphert et al., 2019). Moreover, genetic studies have demonstrated that MafB is dispensable in regulating adult murine islet β cell function, except during pregnancy (Banerjee et al., 2016; Cyphert et al., 2019). MafA expression during mouse β cell development compensates for the absence of MafB, although glucagon secretion from islet α cells is compromised (Conrad et al., 2016). In addition, mis-expressing MafB in *MafA*^*Δpanc*^ β cells cannot rescue MafA function (Cyphert et al., 2019). Collectively, these results illustrate similarities and differences between MafA and MafB in expression and function in rodent islet cells.

While most islet-enriched transcription factors are produced in an analogous fashion in rodents and humans, there are unique differences in the temporal and islet cell type-specific expression pattern of the human MAFA and MAFB proteins. In humans, MAFA protein is not detected in the β cell until about 10 years of age, while MAFB is present throughout the lifespan of human insulin^+^ β cells (Cyphert et al., 2019; Dai et al., 2012). These observations imply that humans have at least two postnatal MAFA^+^ β cell populations capable of maintaining euglycemia, represented by the juvenile (i.e., <10 years old and MAFA^Low^:MAFB^+^) and post-juvenile (≥10+ years old and MAFA^High^:MAFB^+^) periods. In fact, independent studies have established distinct molecular and functional properties of these temporally produced human cell populations (Arda et al., 2016; Arrojo et al., 2019; Camunas-Soler, 2019), and an association between MAFA levels and adult islet β cell functional heterogeneity (Chen et al., 2019). MAFA and MAFB levels are reduced in type 2 diabetic (T2D) islets (Guo et al., 2013), and (at least) MAFA is particularly sensitive to oxidative stress and glucotoxic conditions (Harmon et al., 2005). While the role of MAFA or MAFB has not been analyzed directly in intact human islets, both are required for GSIS in the human EndoC-βH1 β cell line (Scharfmann et al., 2014; Scoville et al., 2015). Moreover, we have recently shown that MAFB is essential to insulin production in human embryonic stem cell derived β cells (Russell et al., 2020), which is in stark contrast to its dispensable role in rodents (Banerjee et al., 2016; Cyphert et al., 2019). These findings highlight species-specific differences in MAFA and MAFB production and islet cell distribution, likely implying that the homodimeric MAFB activator and the MAFA:MAFB heterodimeric activator provide unique functional characteristics to human islet β cells.

Individuals carrying a heterozygous mutation in the MAFA transactivation domain involving a substitution of the conserved Serine at position 64 with a Phenylalanine (S64F MAFA) develop either insulinomatosis, a non-syndromic condition of hyperinsulinemic hypoglycemia caused by multiple insulin-secreting neuroendocrine tumors, or diabetes. The mean age at diagnosis for both conditions is 38 years (Iacovazzo et al., 2018). These results demonstrate that MAFA is a causative gene for Maturity Onset Diabetes of the Young (MODY), a collection of diseases predominantly driven by mutations in essential islet-enriched transcription factors (Barbetti and D’Annunzio, 2018). Interestingly, diabetes is prevalent in male S64F MAFA carriers (i.e., 3:1) while insulinomatosis is more common in female carriers (4:1) (Iacovazzo et al., 2018). *In vitro* analysis demonstrated that the S64F substitution prevents a key priming phosphorylation event at position S65 in MAFA, which profoundly increases the stability of the normally unstable MAFA protein by affecting processes necessary for ubiquitin-mediated degradation (Iacovazzo et al., 2018).

Due to the rare detection of S64F MAFA-mediated diseases and difficulty of performing mechanistic studies in a human context, we used CRISPR/Cas9-based mutagenesis to establish a mouse model (termed *MafA^S64F/+^*) harboring the same pathogenic single base pair substitution (C>T) as human carriers. Diabetes-related phenotypes were specifically associated with male *MafA^S64F/+^* heterozygous mice by 5 weeks of age, which manifested with glucose intolerance and impaired insulin secretion due to reduced glucose-stimulated β cell calcium influx. These changes were preceded by an overt but transient increase in MafA protein levels in islet β cells at 4 weeks. In addition, the functional deficiencies in male *MafA^S64F/+^* islet β cells accompanied the induction of markers of DNA damage, cell cycle exit, senescence, and the senescence-associated secretory phenotype (SASP) at 5 weeks, consistent with accelerated cellular aging and senescence. In contrast, heterozygous mutant female mice not only had lower fasting blood glucose levels and improved glucose clearance, but also did not display any of the overt aging and senescence signatures of littermate, mutant males. Expression of S64F MAFA in the human β cell line, EndoC-βH2, also resulted in increased senescent gene markers and secretion of human-specific SASP factors capable of inducing senescence in a cell non-autonomous manner. Taken together, these results implicate a conserved mechanism involving premature senescence and aging of islet β cells to cause diabetes in human male S64F MAFA carriers.

## Methods

### Animals

S64F MafA-expressing mice were generated using CRISPR/Cas9 targeting at the University of Michigan Transgenic Core in the C57BL/6J mouse strain. Viable animals and progeny were screened for the appropriate mutation by DNA sequencing (Molecular Resource Center, University of Tennessee Health Science Center). Wild-type (WT) littermates were used as controls. *MafA^S64F/+^* were born at normal Mendelian ratios while homozygous *MafA^S64F/S64F^* variants were not (see below). All animal studies were reviewed and approved by the Vanderbilt University Institutional Animal Care and Use Committee. Mice were housed and cared for according to the Vanderbilt Department of Animal Care and the Institutional Animal Care and Use Committee of Animal Welfare Assurance Standards and Guidelines.

### Intraperitoneal glucose and insulin tolerance testing

Glucose tolerance testing was performed on WT and *MafA^S64F/+^* mice (n = 4-16) given an intraperitoneal injection of D-glucose (2 mg/g body weight) prepared in sterile PBS (20% w/v) after a 6-hour fast. Insulin tolerance tests were conducted by intraperitoneal injection of 0.5 IU/kg body weight insulin (Novolin, regular human insulin, recombinant DNA origin) into mice (n=3-5) fasted for 6 hours. Blood glucose was measured using a FreeStyle glucometer (Abbott Diabetes Care) before (0 minutes) and at 15, 30, 60, and 120 minutes following injection. Serum insulin was measured by radioimmunoassay at the Vanderbilt Hormone Assay and Analytical Services Core using blood collected following the 6-hour fast. Serum testosterone was measured by the University of Virginia Center for Research in Reproduction Ligand Assay and Analysis Core.

### Glucose-stimulated hormone secretion

WT and *MafA^S64F/+^* mouse (n = 3-6) islets were isolated using standard islet isolation conditions, and hormone secretion was assessed as described previously (Cyphert et al., 2019). The outcome was presented as the fold change between the percentage of secreted insulin or glucagon relative to the total insulin or glucagon content at either 2.8 or 16.7 mmol/L glucose. Islet insulin and glucagon content was presented as the concentration of insulin or glucagon per DNA content (Quant-iT™ PicoGreen™, Invitrogen) in each reaction (ng/ng DNA).

For perifusion analysis, islets from 5 week old female *MafA^S64F/+^* and controls were studied in a dynamic cell perifusion system at a perifusate flow rate of 1 mL/min (Walker, 2020) by the Vanderbilt Islet Procurement and Analysis Core. The effluent was collected at 3-minute intervals using an automatic fraction collector. The insulin concentration in each perifusion fraction and islet extract was measured by radioimmunoassay (Millipore).

### Tissue preparation and immunostaining

For immunostaining, WT and *MafA^S64F/+^* pancreata were fixed in 4% (v/v) paraformaldehyde, embedded in either Tissue-Plus O.C.T. (Thermo Fisher Scientific) or paraffin wax, and sectioned to 6 μm thickness. Immunofluorescent images were obtained using the Zeiss Axio Imager M2 widefield microscope with ApoTome. Immunofluorescence staining was performed as previously described with the antibodies listed in **Supplementary Table 1**. Islet β- and α- cell areas were determined as described previously (Cyphert et al., 2019). Briefly, pancreatic sections taken every 50μm (n=3-4 animals per genotype) were scanned using a ScanScope CS scanner (Aperio Technologies, Vista, CA). Images from each experiment were processed with ImageScope Software (Aperio Technologies, Vista, CA). Islet β- and α-cell areas were calculated as the ratio of the insulin- or glucagon-positive area to total pancreas area (eosin stained). The TUNEL assay was performed using an *in situ* cell death detection kit (Roche, #11684795910).

### RNA sequencing and analysis

RNA was isolated from WT and *MafA^S64F/+^* islets (n=4 for males, n=5 for females) using the RNAqueous total RNA isolation kit (Ambion; Thermo Fisher), and then analyzed on an Agilent 2100 Bioanalyzer. Only samples with an RNA Integrity Number >8.0 were used for library preparation. cDNA libraries were constructed and paired-end sequencing of 4-5 replicates was performed on an Illumina NovaSeq6000 (150 nucleotide reads). The generated FASTQ files were processed and interpreted using the Genialis visual informatics platform (https://www.genialis.com). Sequence quality checks were determined using raw and trimmed reads with FastQC (http://www.bioinformatics.babraham.ac.uk/projects/fastqc), and Trimmomatic (Bolger et al., 2014) was used to trim adapters and filter out poor-quality reads. Trimmed reads were then mapped to the University of California, Santa Cruz, mm10 reference genome using the HISAT2 aligner (Kim et al., 2015). Gene expression levels were quantified with HTSeq-count (Anders et al., 2015), and differential gene expression analyses performed with DESeq2 (Love et al., 2014). Poorly expressed genes, which had expression count summed over all samples of <10, were filtered out from the differential expression analysis input matrix. Quantitative real-time PCR expression analysis of selected candidates was performed using cDNAs produced from the iScript cDNA Synthesis Kit (Bio-Rad) in a LightCycler 480 system (Roche) with primers provided in **Supplementary Table 2**.

### Islet cytosolic calcium imaging

Approximately 60 WT and *MafA^S64F/+^* islets from at least 6 female and male mice were analyzed with the ratiometric calcium indicator fura-2-acetoxymethylester (Fura-2 AM) (Life Technologies). Islets were maintained in 5 mM glucose for 30 min prior to measuring 11 mM glucose-induced calcium oscillations and depolarization-activated calcium influx with 30 mM KCl. Islets were loaded with 2 μM Fura-2 AM for 20 min, washed, transferred to a hand-made agarose gel small-volume chamber in a glass bottom dish. Images were taken using a Nikon Eclipse TE2000-U microscope equipped with an epifluorescence illuminator (SUTTER, Inc), a CCD camera (HQ2; Photometrics, Inc), and Nikon Elements software (NIKON, Inc) as described before (Dadi et al., 2015).

### Senescence-associated β-galactosidase (SA-β-gal) staining

WT and *MafA^S64F/+^* pancreata were snap frozen in O.C.T. and cryosections were prepared at 16 μm thickness. SA-β-gal activity staining was performed at pH 6.0 (Kurz et al., 2000) using a commercial kit (Cell Signaling, #9860). To compare the intensity of SA-β-gal staining, sections from different genotypes and ages were processed on the same slide. Staining reactions were developed for 18 hours at 37°C, then stopped by 3x PBS washes (pH 7.4). Slides were then subject to immunostaining for insulin by fixing in 4% paraformaldehyde for 45 minutes, permeabilized with Tris-buffered saline with 0.2% Triton-X-100 for 15 minutes, blocked in 2% normal donkey serum/1% BSA in PBS and incubated overnight with guinea pig anti-insulin (1:500, Abcam) at 4°C. HRP-conjugated secondary antibodies were then incubated on slides for 1-hour and detected with the DAB+ chromogen kit (DAKO). After washing, slides were mounted and imaged by brightfield microscopy. The total number of SA-β-gal^+^ islets per pancreas section were analyzed on ImageJ.

### Ovariectomy methods

WT and *MafA^S64F/+^* female mice underwent ovariectomy at 3 weeks to remove the contribution of endogenous ovarian hormones and prevent completion of sexual maturity (Martinez et al., 2012). Mice were anesthetized with 1-5% inhaled isoflurane, and placed prone on a heating pad to maintain their body temperature at 37° C. The mice received subcutaneous ketofen at 5-10 mg/kg prior to surgery and post-operatively for two days for pain relief. Intraperitoneal ceftriaxone at 20-40 mg/kg was given once intraoperatively for infection prophylaxis. After adequate anesthesia and analgesia, a midline 1.5 cm incision was made along the shaved mid-dorsal surface of the mouse. Within the retroperitoneal cavity, the ovaries were located and ligated. The incision was closed using surgical staples, which were removed on post-operative day three. Mice were housed singly during recovery and regrouped post-operatively.

### Glycogen storage assay

Liver and gastrocnemius muscle from WT and *MafA^S64F/+^* mice (n=5) were collected and flash frozen in liquid nitrogen. Tissue was homogenized with a Dounce homogenizer prior to determining glycogen content using a Glycogen assay kit (Cayman Chemical) according to the manufacturer’s protocol.

### Human EndoC-βH2 cells

Human EndoC-βH2 cells were grown in DMEM containing 5.6mM glucose, 2% BSA,50 μM 2-mercaptoethanol, 10mM nicotinamide, 5.5 μg/mL transferrin, 6.7 ng/mL selenite, 100 units/mL penicillin, and 100 units/mL streptomycin (Scharfmann et al., 2014). For gene transfection, cells were incubated with lentiviral particles (150ng for cells cultured in a 6cm dish) containing either WT MAFA-expressing or S64F MAFA-expressing sequences one day after plating. SA-β-gal staining was performed at pH 6.0 (Kurz et al., 2000) using a commercial kit (Cell Signaling, #9860). To compare the intensity of SA-β-gal staining, cells from different conditions were cultured on chamber slides and processed on the same slide. Staining reactions were stopped after developing for 2-3-hours at 37°C with 3x PBS washes (pH 7.4). Slides were then subject to immunostaining for MAFA by fixing in 4% paraformaldehyde for 12 minutes, permeabilized with Tris-buffered saline with 0.5% Triton-X-100 for 5 minutes, blocked in 2% normal donkey serum/1% BSA in PBS and incubated overnight with Rabbit α-MAFA (1:500, Novus (NBP1-00121) at 4°C. After washing, slides were mounted and imaged on a Zeiss Axio Imager M2 widefield microscope with ApoTome. SA-β-gal+ cell numbers were analyzed on ImageJ. RNA was collected using Trizol reagent (Life Technologies) 1 week following infection. The iScript cDNA synthesis kit (Bio-Rad Laboratories, Inc.) was used for cDNA synthesis. The quantitative (q)PCR reactions were performed on a LightCycler 480 II (Roche), and analyzed by the ΔΔCT method with primers provided in **Supplementary Table 2**. Significance was calculated by comparing the ΔCT values.

### Paracrine Conditioned Media Assay

Conditioned media (CM) produced after 24 hours of culture from S64F MAFA and WT MAFA-expressing human EndoC-βH2 cells was clarified by centrifugation at 500xg for 5 minutes and then at 3000xg for 5 minutes. CM was stored at 4°C for up to 3 weeks prior to use. EndoC-βH2 cells plated on 12-well plate were cultured with a 1:1 mix of EndoC-βH2 islet media (4% BSA) and albumin-free CM collected from WT MAFA or S64F MAFA expressing cells (resulting in 2% BSA in the final mix). For controls, EndoC-βH2 cells from the same passage were cultured in a 1:1 mix of regular EndoC-βH2 cells media and albumin-free media alone (not conditioned by cells), also resulting in 2% BSA in the final mix. After 72 hours, EndoC-βH2 cells were harvested for qRT-PCR as described above.

### Statistical Analysis

Statistical significance was determined using the two-tailed Student *t* test. Data are presented as the mean ± SEM. A threshold of *P* < 0.05 was used to declare significance.

## Results

### Male *MafA*^*S64F/+*^ mice are glucose intolerant due to impaired insulin secretion

Since MafA is a primary regulator of glucose clearance and insulin secretion in islet β cells (Hang and Stein, 2011), *MafA^S64F/+^* and WT littermates were subjected to glucose tolerance testing (GTT) and fasting blood glucose measurements at various postnatal time points. While both male and female heterozygous animals had improved glucose clearance at 4 weeks, females also had modest fasting hypoglycemia (**Figure 1A**). Strikingly, male heterozygous mutant animals developed persistent glucose intolerance beginning at 5 weeks of age, while females continued to have improved glucose tolerance and lower fasting blood glucose levels (**Figure 1A**). GTT results were stably maintained in male and female *MafA^S64F/+^* mice (**Supplemental Figure 1**). The sex-dependent male phenotype was even more penetrant in homozygous *MafA^S64F/S64F^* mutants, as males displayed overt diabetes with elevated fasting glucose levels that progressively worsened (**Supplemental Figure 2A**, **B**). In contrast, the phenotype of homozygous females was much more variable and largely comparable to WT littermates (**Supplemental Figure 2C**). Because of the generally poor survival of homozygous mutants postnatally (**Supplemental Figure 2D**), possibly resulting from severe neurodevelopmental defects due to *MafA^S64F/S64F^* expression and its impaired activity in the dorsal root ganglia (Lecoin et al., 2010; Niceta et al., 2015), all of the remaining experimentation was performed with male and female *MafA^S64F/+^* mice.

**Figure 1:**
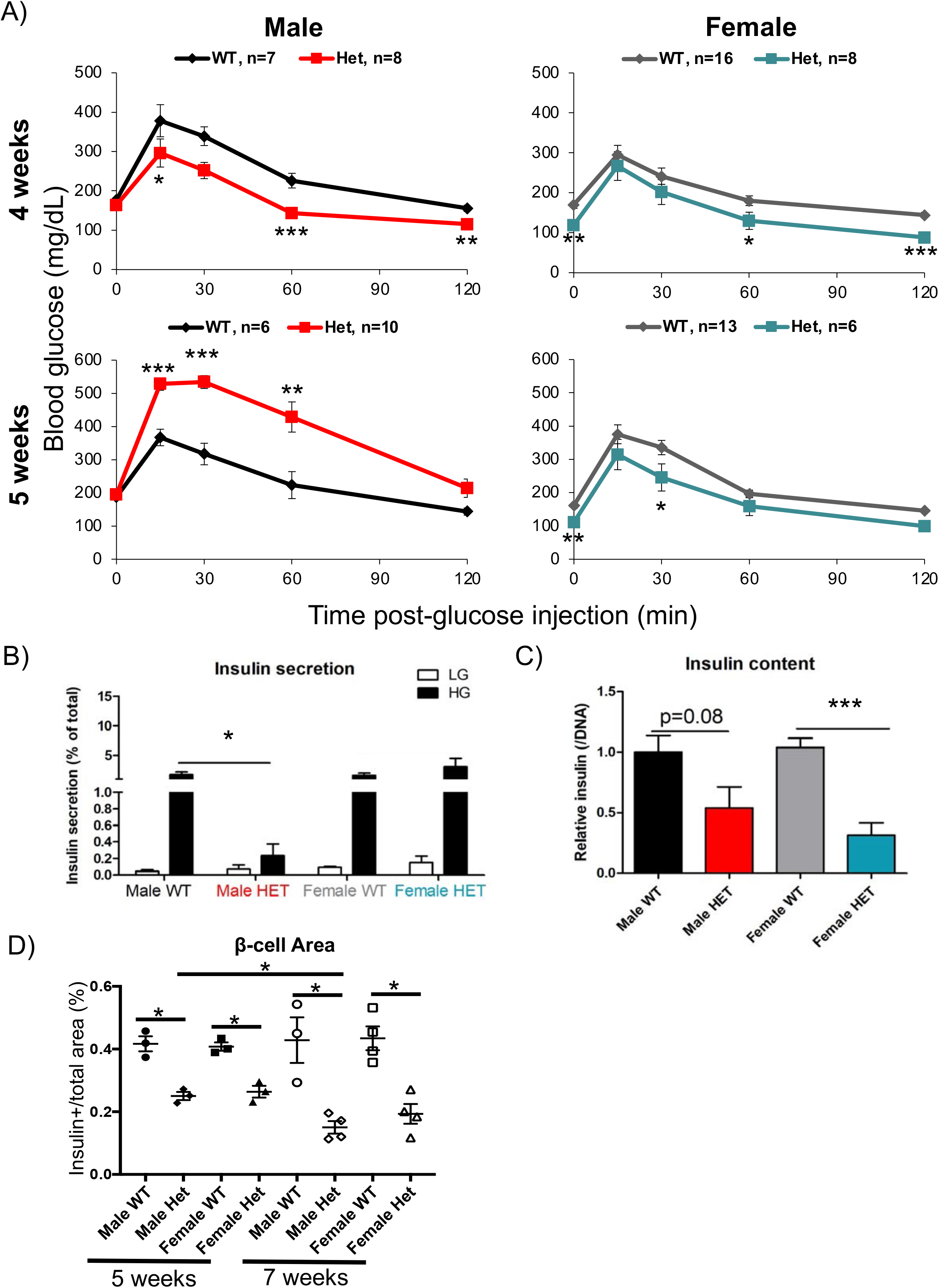
Male but not female *MafA^S64F/+^* mice become glucose intolerant between 4 and 5 weeks of age. A) Fasted male and female animals underwent intraperitoneal glucose tolerance tests at 4 and 5 weeks of age. Male heterozygous (termed Het) *MafA^S64F/+^* mice (red line) had improved glucose clearance at 4 weeks but become glucose intolerant by 5 weeks. Female Het mice (teal line) had significantly lower fasting blood glucose levels and improved glucose clearance at both time points. B) High glucose stimulated insulin secretion in isolated islets was impaired in male Het samples at 5 weeks. Islets were incubated with 4.6 mM (low, LG) or 16.8 mM (high, HG) glucose for 1-hour. C) Islet insulin content trended lower in Het males and was significantly decreased in females. Levels were normalized to insulin content (in panel B) and DNA content (in panel C). D) Male and female islet β cell area was reduced at 5 weeks in *MafA^S64F/+^* mice and even further by 7 weeks. The area was calculated by dividing the total Insulin^+^ area by the total pancreatic area (eosin staining) in pancreas sections obtained every 50 μm multiplied by 100 to obtain percent (%). *p<0.05; **p<0.01; ***p<0.001.

*Ex vivo* GSIS was impaired in 5-week-old male *MafA^S64F/+^* islet β cells (**Figure 1B**), although insulin content trended lower in male islets and was significantly reduced in female islets (**Figure 1C**). *MafA^S64F/+^* female mice appear to have lower fasting blood glucose levels (**Figure 1A**) due to increased insulin secretion in both low (5.6) and high (16.7) glucose conditions as determined by dynamic perifusion assays (**Supplemental Figure 3A**) and increased fasting serum insulin levels (**Supplemental Figure 3B**). Because of the counter-regulatory effects of insulin on glucagon hormone secretion (Franklin et al., 2005), we considered that glucose-regulated glucagon levels might be altered in *MafA^S64F/+^* mice. However, there was neither an obvious sex-dependent impact on glucose-stimulated glucagon secretion nor a change in glucagon content (**Supplemental Figure 4A**, **B**). Islet α cell area was decreased in both male and female heterozygous islets, although this was variable among animals (**Supplemental Figure 4C**).

Insulin^+^ β cell area was reduced in *MafA^S64F/+^* mice of both sexes (**Figure 1D**), which may be due to their lower cell proliferation rates in *MafA^S64F/+^* mice relative to WT littermates (**Supplemental Figure 5**). In addition, the changes in blood glucose levels were not attributable to differences in peripheral tissue insulin sensitivity or glucose uptake as insulin tolerance tests and glycogen storage were both unchanged (**Supplemental Figure 6A, B**). Animal growth rates were also not appreciably altered as assessed by body weight (**Supplemental Figure 6C**).

Because the sex hormones estradiol and testosterone impact islet cell function (Gannon et al., 2018), we questioned whether these hormones influenced the sex-biased phenotypes in *MafA^S64F/+^* mice. Both sexes achieved puberty appropriately with fertility comparable to WT littermates (data not shown). Female *MafA^S64F/+^* mice were ovariectomized to investigate whether estrogen was regulating mutant sex-biased activity. Ovariectomy at 3 weeks of age produced glucose intolerance in WT female mice by 4 weeks of age. In contrast, glucose tolerance was unaffected by ovariectomy in age-matched female *MafA^S64F/+^* mice (**Supplemental Figure 7A,** data not shown). These results indicate that the S64F MafA mutation is dominant over the effect of estrogen-deficiency on glucose intolerance. Furthermore, testosterone levels were not altered markedly in males at both 4 and 5 weeks of age and were similar to age-matched females in these pre- and peri-pubertal mice (**Supplemental Figure 7B**). The low circulating levels of testosterone produced during this phenotypic window lowers the likelihood of this hormone influencing MafA^S64F^ actions.

Collectively, these data strongly suggested that S64F MafA regulates male and female islet β cells through common and distinct mechanisms; however, these processes are not principally influenced by the primary sex hormones.

### MafA protein levels are transiently and profoundly upregulated prior to the changes in glucose clearance in male *MafA*^*S64F/+*^ islet β cells

The S64F MAFA variant impairs a key, priming phosphorylation event at position S65, which blocks subsequent GSK3-mediated phosphorylation at positions S61, T51, T57, and S49 in the transactivation domain. These changes impede ubiquitin-mediated protein degradation of S64F MAFA, and its stability is dramatically increased in the human EndoC-βH1 β cell line (Han et al., 2007; Iacovazzo et al., 2018; Rocques et al., 2007). Consequently, we predicted that this mutation in the highly conserved region of the protein (Han et al., 2007; Iacovazzo et al., 2018; Rocques et al., 2007) would also increase MafA levels in *MafA^S64F/+^* β cells. Surprisingly, we only found elevated MafA protein immunostaining intensity in male *MafA^S64F/+^* islets at 4 weeks of age (**Figure 2A**), one week prior to the onset of glucose intolerance (**Figure 1A**). In contrast, MafA protein levels were not increased in female *MafA^S64F/+^* β cells at 4 weeks or any other analyzed time point in relation to WT controls (**Figure 2B**). However, there was a roughly 3- to 5-fold decrease in *MafA* transcript levels at 4 and 5 weeks of age in both male and female *MafA^S64F/+^* islets compared to WT littermates (**Figure 2C**). The reduction in *MafA* gene expression indicates that the more stable MafA^S64F^ protein is acting in an autoregulatory manner by binding at the conserved MafA binding site within the 5’-flanking sequences of the Region 3 transcription control domain (Artner et al., 2008; Artner et al., 2010; Scoville et al., 2015).

**Figure 2:**
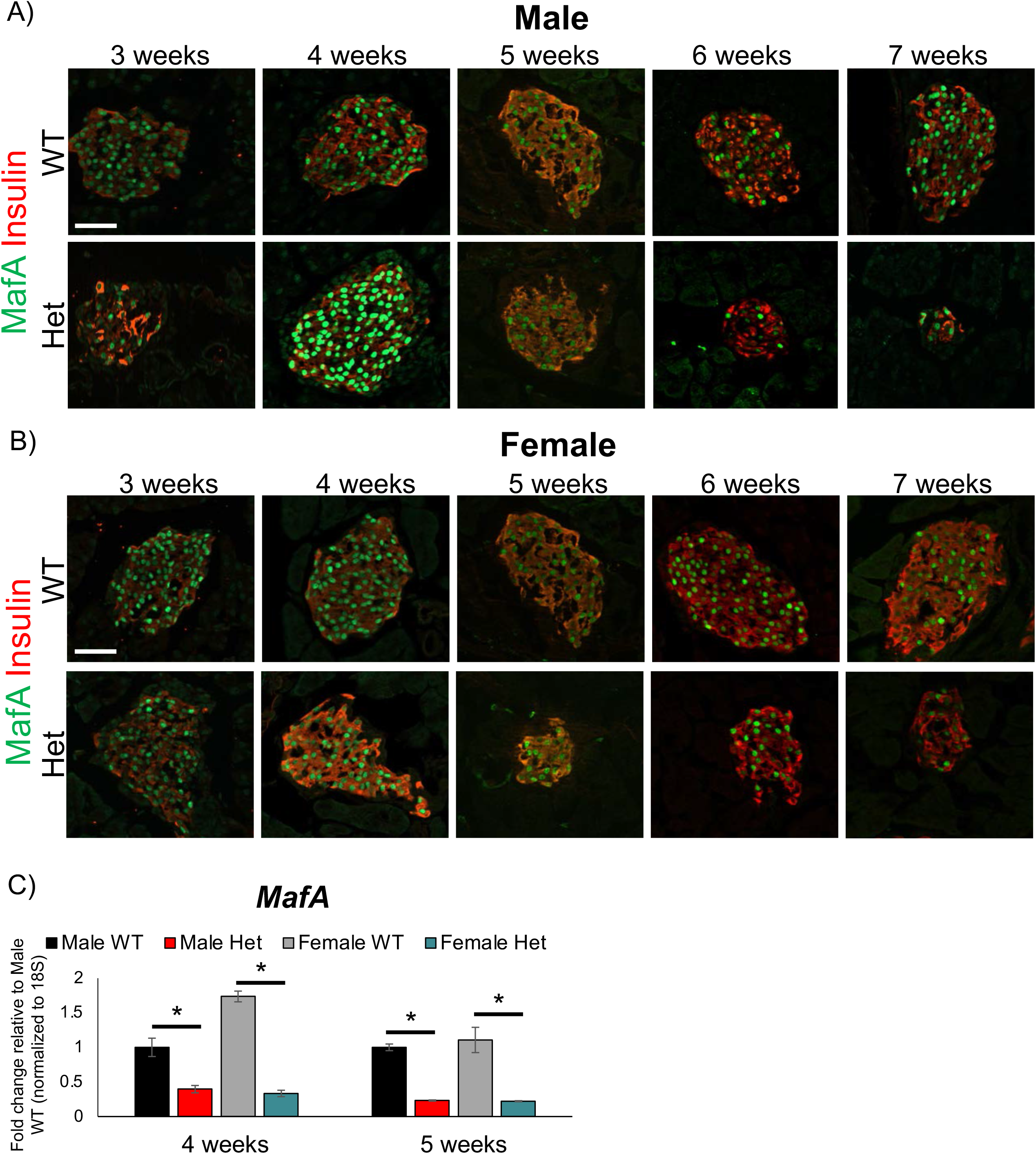
MAFA protein levels were transiently upregulated at 4 weeks in male *MafA^S64F/+^* islets. A) Immunostaining over the course of 3, 4, 5, 6 and 7 weeks revealed that MAFA protein staining intensity only increased at 4 weeks of age in male *MafA^S64F/+^* (Het) islets. B) In contrast, MAFA staining intensity was unchanged between female Het and WT islets. C) *MafA* mRNA levels were significantly reduced at 4 and 5 weeks in both male and female Het islet samples. Fold change shown relative to male WT islets. *p<0.05; scale bars=50μm.

### Glucose- and potassium chloride (KCl)-stimulated calcium handling were altered in both male and female *MafA*^*S64F/+*^ islets

To provide an unbiased and comprehensive perspective on how MafA^S64F^ influences islet β cell gene expression, bulk RNA-sequencing was performed on isolated islets from 5-week-old WT and *MafA^S64F/+^* male and female mice when a clear delineation of phenotypes was observed. Female heterozygous islets had 736 differentially expressed (DE) genes (337 up and 399 down) compared to female WT islets, while males had 2410 DE genes (2031 up and 379 down) compared with male WT islets (**Figure 3A**). There were also 391 DE genes that were similarly regulated between sexes, which were revealed by gene ontology analysis to include factors associated with Ca^+2^ and K^+1^ channels important in controlling β cell function (**Figure 3B, C**). In addition, other voltage-dependent calcium channels genes as well as those genes important to ion influx were also specifically up-regulated in male *MafA^S64F/+^* islets (**Figure 3D**), suggesting both sex-dependent and -independent changes in ion flux in *MafA^S64F/+^* islets.

**Figure 3:**
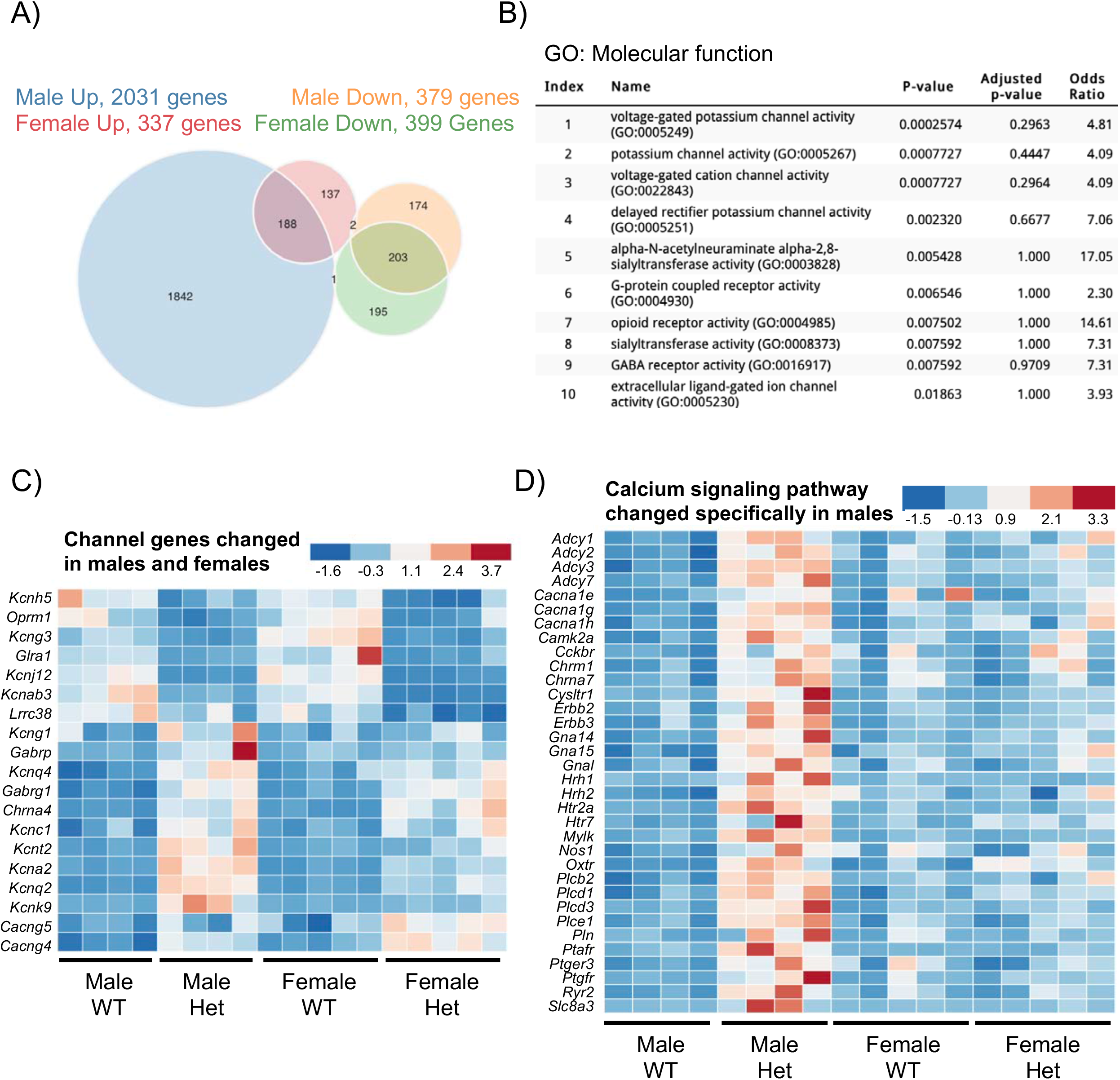
Male and female *MafA^S64F/+^* islet cells regulate a common and distinct set of genes associated with Ca^+2^ and K^+1^ channel activity. A) The Venn diagram illustrates the total number of RNA-Seq identified genes differentially up- or down-regulated between 5-week-old WT and Het islets. B) Gene Ontology (GO): Molecular function analysis of the 391 genes commonly up- or down-regulated in *MafA^S64F/+^* (Het) islets revealed alterations in multiple ion channel activity pathways. C) Heat maps showing channel gene expression changes common between male and female Het islets, and (D) Ca^+2^ signaling pathway genes uniquely increased in male Het islets by KEGG analysis (also see Figure 5A). FDR<0.05.

Since glucose-induced elevations of cytoplasmic Ca^+2^ result in insulin release (Bergsten et al., 1994), *MafA^+/+^* and *MafA^S64F/+^* islet Ca^+2^ handling was monitored in response to glucose stimulation and KCl-induced depolarization. As expected from the diminished GSIS response of male *MafA^S64F/+^* islets (**Figure 1B**), their Ca^+2^ entry was significantly blunted relative to the WT islets in response to 11 mM glucose stimulation (**Figure 4A**; represented by red trace in left panel and termed “non-responder”). The glucose-stimulated calcium oscillation pattern that is responsible for pulsatile insulin secretion was also significantly diminished in *MafA^S64F/+^* islets (**Figure 4A**). In contrast, there appeared to be two distinct islet populations in female *MafA^S64F/+^* animals with different glucose-stimulated Ca^+2^ responses. One was similar to male *MafA^S64F/+^* islets in having a very limited initial glucose response and minimal oscillatory behavior (i.e., **Figure 4A,** dark green representative trace in right panel and labeled as “non-responder”). However, the remaining islets had more subtle decreases in the initial response to 11 mM glucose, and subsequently oscillate with higher frequency but equivalent amplitude to WT islets (**Figure 4A;**“responders”, teal representative trace in right panel). The average peak amplitude of the initial glucose-induced calcium response was reduced in all male and female *MafA^S64F/+^* islets (**Figure 4B**). This suggests that the *MafA^S64F/+^* female islet “responders” population is able to compensate for the “non-responders” and maintain glucose tolerance in *MafA^S64F/+^* female mice, which would be predicted because only a fraction (roughly 20%) of the β cell mass is thought to be required to maintain glucose tolerance (Gale, 2002).

**Figure 4:**
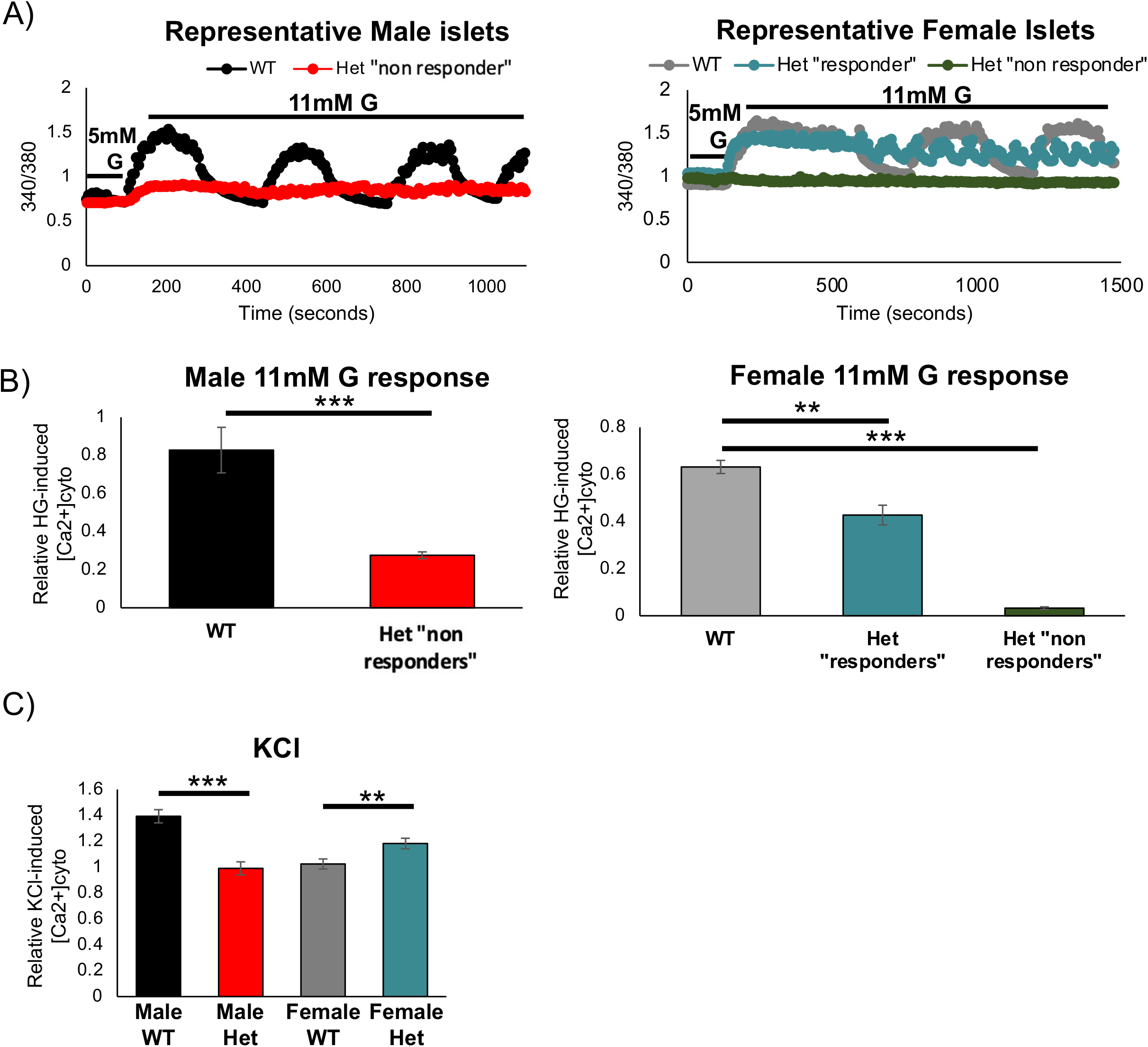
Glucose-induced Ca^+2^ oscillations and KCl-induced Ca^+2^ responses are altered in *MafA^S64F/+^* islets. A) Representative Fura2 traces show loss of the β cell-induced Ca^+2^ oscillations by 11 mM glucose (G) in 5-week-old male *MafA^S64F/+^* (Het) islets (red line and termed non-responders). Female Het islets had two different functionally responsive islet populations (“responders,” teal line; “non-responders,” green line). B) Quantitation of male (left) and female (right) islets showed reduced cytoplasmic Ca^+2^ following stimulation with 11 mM glucose. The average peak amplitude was quantitated by dividing the first peak Δcalcium after 11mM glucose application by the baseline response at 5mM glucose. C) The Ca^+2^ response to 30mM KCl is reduced in male Het islets but increased in female Het islets. Representative traces for this experiment are shown in Supplemental Figure 8. **p<0.01; ***p<0.001.

Male and female *MafA^S64F/+^* islets also had disparate Ca^+2^ responses to KCl-mediated depolarization. The KCl-induced calcium influx (Δcalcium after KCl application divided by baseline calcium) was significantly reduced in male *MafA^S64F/+^* islets when compared to WT male islets (**Figure 4C** and **Supplemental Figure 8A**). Interestingly, however, the female *MafA^S64F/+^* islets showed greater KCl-induced calcium influx when compared to WT female islets, which was observed for both glucose-stimulated Ca^+2^ “responders” and “non-responders” (**Figure 4C**, **Supplemental Figures 3A** and **8A**). Importantly, female *MafA^S64F/+^* islets also showed greater baseline (5 mM glucose) Ca^+2^ signaling than WT female (**Supplemental Figure 8B**), which is consistent with the fasting hypoglycemia and increased fasting insulin levels observed in female *MafA^S64F/+^* mice (**Figure 1A** and **Supplemental Figure 3B**).

### Only male *MafA*^*S64F/+*^ islets express markers of accelerated cellular senescence and aging

Because 5-week-old male *MafA^S64F/+^* islet β cells were not only defined by a predominant, calcium non-responsive population (**Figure 4A, B**) but also by glucose intolerance (**Figure 1A**), we focused our attention on identifying molecular determinants to their poor function. In addition to the previously illustrated gene expression alterations involved in calcium signaling (Index ranking #6, **Figure 5A**), 5-week-old male *MafA^S64F/+^* RNA-Seq data revealed upregulation of many metabolic pathway genes implicated in cellular aging (e.g. Index ranking pathways #4, 6, and 8) and senescence (e.g. Index ranking pathways #1, 2, 3, 5, 6, 7 and 8 in **Figure 5A**) (Basisty et al., 2018; Campisi and d’Adda di Fagagna, 2007; Coppe et al., 2010; Franceschi et al., 2018; Greenhill, 2019; Kabir et al., 2016; Mancini et al., 2012; Martin and Bernard, 2018). Furthermore, the preponderance of upregulated genes detected by RNA-Seq indicates that MafA^S64F^ is acting as a dominant activator in (at least) male *MafA^S64F/+^* islet β cells (**Figure 3A**).

**Figure 5:**
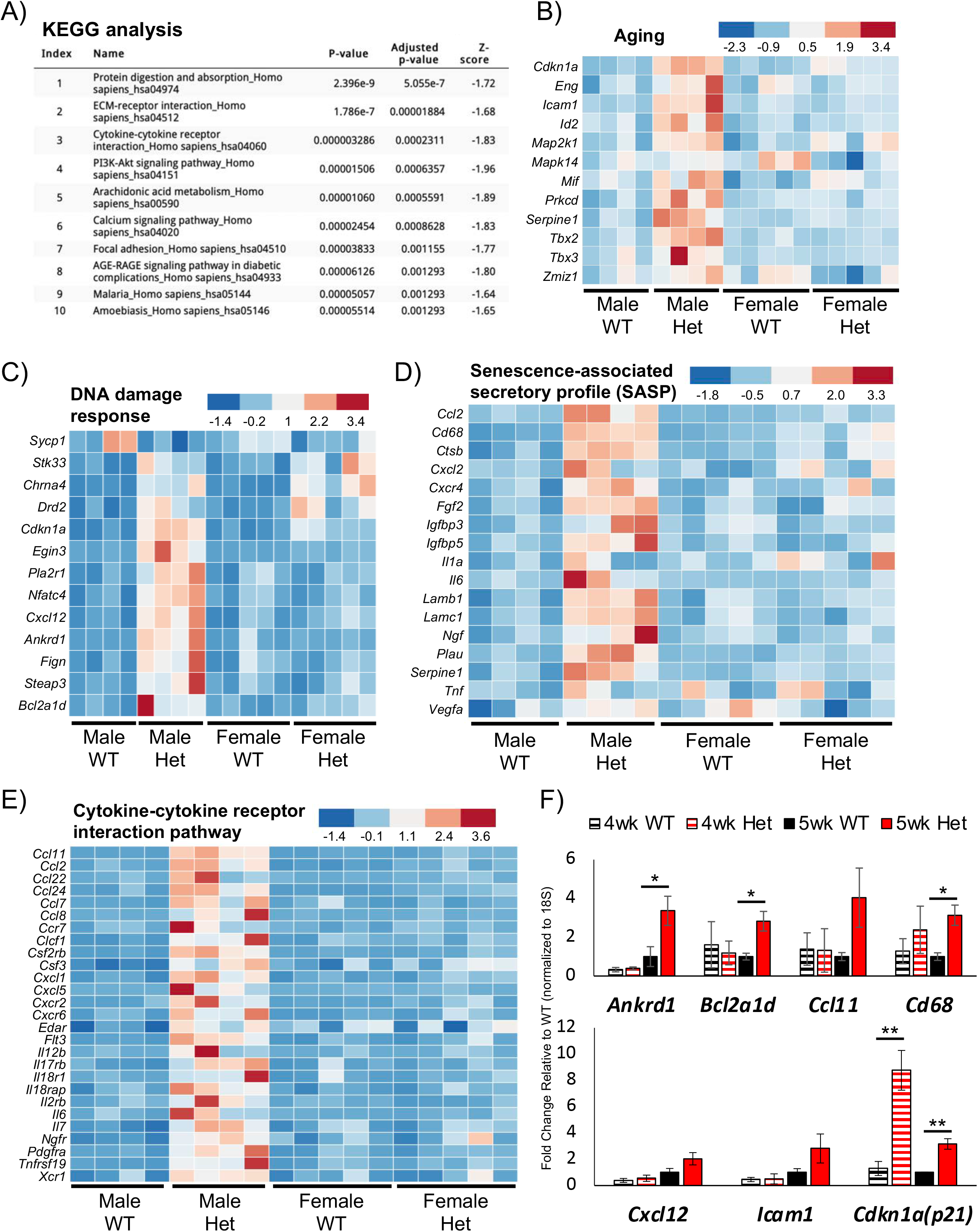
Male *MafA^S64F/+^* islets display increased expression of aging, DDR, SASP, and cytokine pathway gene signatures. A) KEGG analysis of the 1842 genes specifically upregulated in 5-week-old male *MafA^S64F/+^* (Het) islets. Heat maps reveal male Het islets have increased expression of pathway genes associated with B) aging, C) DDR, D) SASP, and E) cytokine-cytokine receptor interactions. FDR<0.05. F) qRT-PCR confirmation of pathway gene changes in 5-week-old male Het islets (solid bars). Gene expression was unchanged at 4-week-old male Het islets with the exception of Cdkn1a (p21) upregulation (hashed bars). *p<0.05; **p<0.01.

Cellular senescence is a durable, cell cycle arrest response that regulates cell fate specification, tissue patterning, and function during development, cell maturation, and organismal aging (e.g., islet β cells (Campisi, 2014; Gorgoulis et al., 2019; Helman et al., 2016; Herranz and Gil, 2018)). It is also induced in a variety of disease states and primarily serves as a stress response to many internal and external insults (Gorgoulis et al., 2019; Herranz and Gil, 2018). The importance of premature aging and senescence in driving male *MafA^S64F/+^* β cell dysfunction was supported by the many genes associated with these responses, including those controlling Ca^+2^ signaling (**Figure 3D**), aging (**Figure 5B**), DNA damage response (**DDR**, **Figure 5C**), the senescence-associated secretory phenotype (**SASP**, **Figure 5D**), and cytokine-cytokine receptor interactions (**Figure 5E**). As expected, the expression of candidate genes linked to aging and senescence were increased in 5-week-old male *MafA^S64F/+^* islets (**Figures 5F**), although not prior to manifesting glucose intolerance in 4-week-old males (**Figures 1A and 5F**). Importantly, these genes were unaltered in *MafA^S64F/+^* female islets (**Supplemental Figure 9A**), although many female differentially expressed genes were also verifiable by qPCR (**Figures 3D**, **5B-F** and **Supplemental Figure 9B, C**).

Senescent cells undergo a progression of changes after an initial insult: early initiation of cellular arrest by activation of cell cycle inhibitors (e.g., p53, p21 and/or p16) and DDR responses in an attempt to regain homeostasis (Gorgoulis et al., 2019; Herranz and Gil, 2018). DDR can be characterized by markers such as phosphorylated histone H2AX (gH2AX) and 53BP1 recruitment to chromatin. We identified increased immunostaining for the γH2AX marker of DNA double-strand breaks, the P53 binding protein-1 (53BP1) DNA damage marker (Schultz et al., 2000), and the p21 cell cycle arrest and senescence marker (Fang et al., 1999) in 5-week-old *MafA^S64F/+^* male, but not female islets (**Figure 6A, B, C** and **Supplemental Figure 10A, B, C**). Indeed, the proportion of islet β cells with p21^+^ staining greater than 10% increased significantly in male *MafA^S64F/+^* islets (**Figure 6C**, right panel).

**Figure 6:**
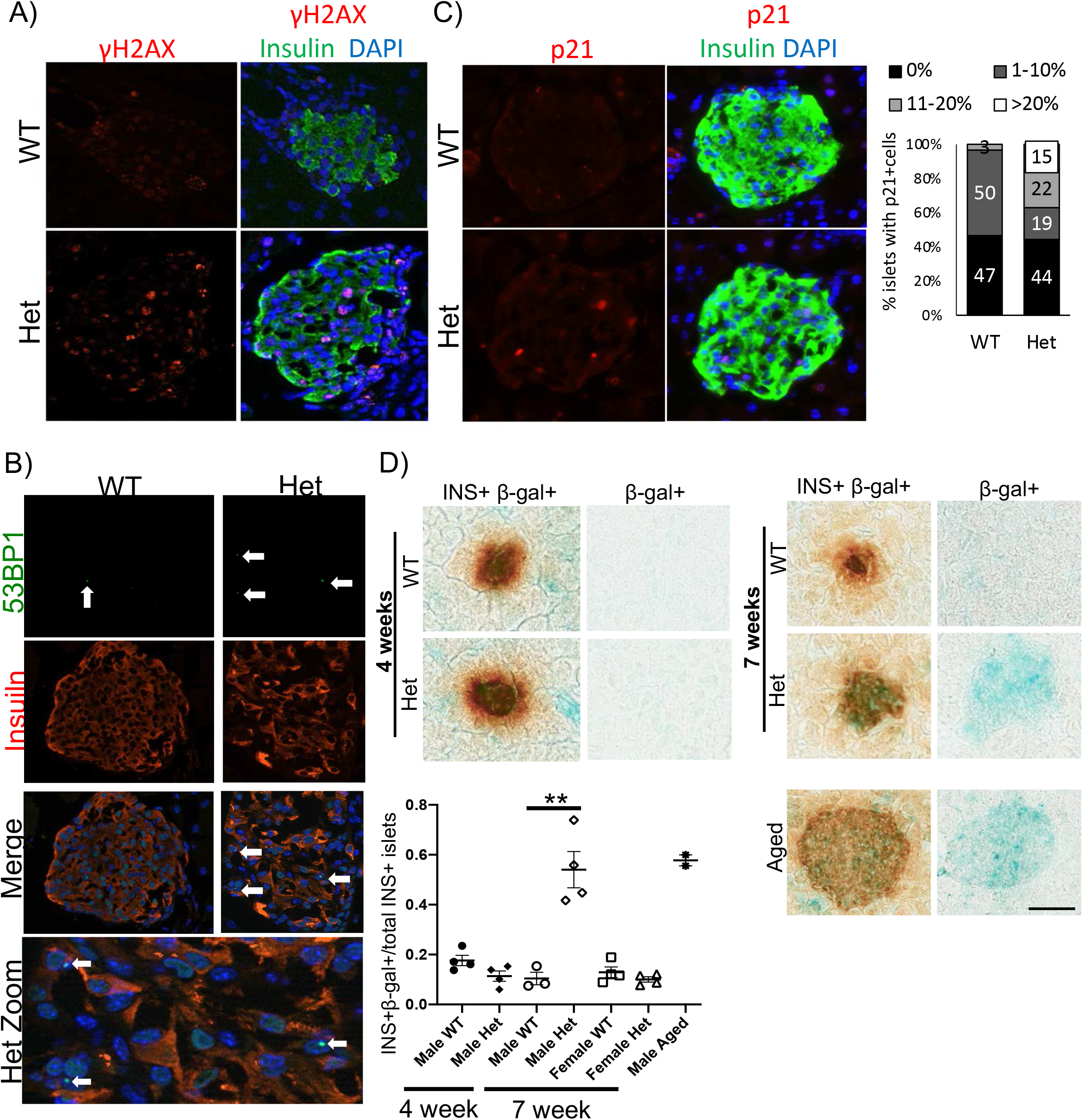
Senescence markers are increased in dysfunctional male *MafA^S64F/+^* islets. γH2AX staining (A) and 53BP1 (B), markers of DNA double-strand breaks, and p21, a cell cycle inhibitor (C), were present in male *MafA^S64F/+^* (Het) islets. Male Het islets had a significant increase in the proportion of islets with >10% p21^+^ cells. D) SA-β-gal was not produced in 4-week-old male Het islets but was detected in Het males by 7 weeks. The SA-β-gal in 7-week Het islets was of similar intensity to 10-12 month-old WT male mouse islets. The proportion of SA-β-gal^+^ islets was quantitated for each sample (n=3-4). **p<0.01; scale bar=50μm.

Apoptosis was not induced in response to DNA damage in either male or female *MafA^S64F/+^* islets, as concluded by the inability to detect islet TUNEL^+^ cells between 4-7 weeks of age (**Supplemental Figure 11,** data not shown). Indeed, induction of anti-apoptosis genes (such as the BCL2 family members; **Figure 5F**) and resistance to apoptosis are consistent with the senescence phenotype (Herranz and Gil, 2018; Thompson et al., 2019). In fact, increased endogenous senescence-associated β-galactosidase (SA-β-gal) staining was detected in male *MafA^S64F/+^* islets at levels comparable to aged mice (**Figure 6D**). Notably, SA-β-gal was undetectable in male *MafA^S64F/+^* islets prior to compromised β cell function at 4 weeks or in female mutant islets (**Figure 6D** and **Supplemental Figure 10D**). Together, these results illustrate a novel sex-dependent pathophysiology of accelerated β cell aging and senescence that contributes impaired GSIS and glucose intolerance in *MafA^S64F/+^* males.

### Human EndoC-βH2 β cells expressing MAFA^S64F^ show increased markers of cellular senescence and a functional senescence-associated secretory phenotype

We transduced human EndoC-βH2 cells with human WT MAFA or MAFA^S64F^ expression constructs using lentivirus to investigate whether cellular senescence could be prematurely induced in human β cells. As in our *MafA^S64F/+^* mouse model, MAFA^S64F^ producing cells had significantly increased SA-β-gal expression compared to those expressing the human WT protein (**Figure 7A, B**). Immunoblot analysis of protein extracts from transduced EndoC-βH2 cells also confirmed the faster migrating properties of the phosphorylation impaired MAFA^S64F^ (**Figure 7C**) (Iacovazzo et al., 2018). Gene expression analysis of MAFA^S64F^ cells showed increases in a subset of senescence markers identified in male *MafA^S64F/+^* mice (*P21, BCL2*) and decreased expression of *MAFB* (**Figure 7D**), a transcription factor not produced in rodent islet β cells postnatally (Conrad et al., 2016).

**Figure 7.**
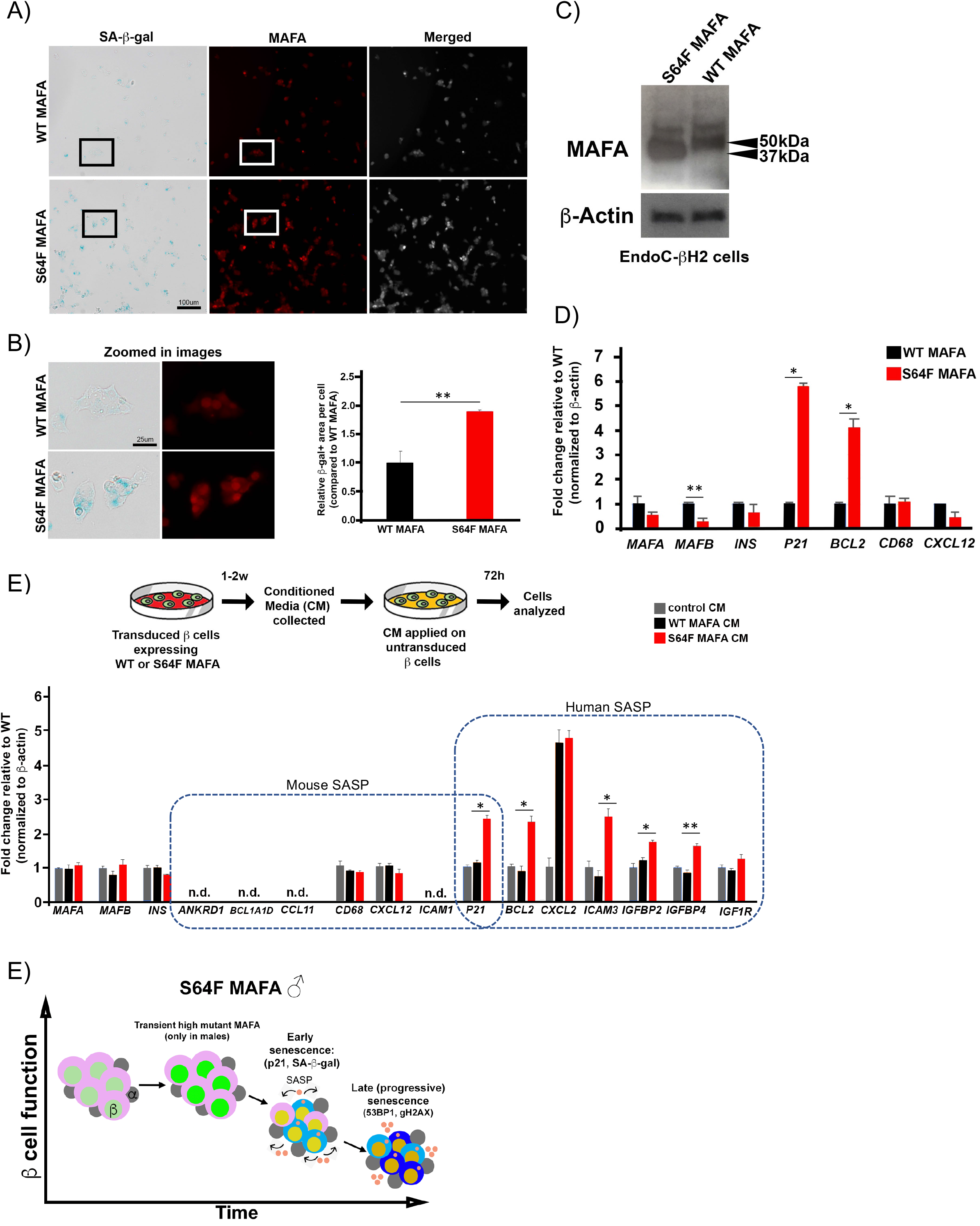
Human β cells expressing MAFA^S64F^ demonstrate accelerated senescence and a species-specific SASP signature. A) SA-β-gal staining on EndoC-βH2 cells transduced to express WT MAFA or MAFA^S64F^ and stained for MAFA (red). Scale bar=100μm. B) Inset areas outlined in boxes in (A) are magnified in (B, left panel). Scale bar=25μm. β-gal^+^ area was significantly increased in MAFA^S64F^ cells (B, right panel). C) Western blotting for EndoC-βH2 cells transduced to express WT MAFA or MAFA^S64F^ shows faster migration in the MAFA^S64F^ lane consistent with under-phosphorylated MAFA. Increased MAFA protein stability is notable in MAFA^S64F^-expressing cells, similar to that seen in (Iacovazzo et al., 2018). D) Human expression of β cell-enriched proteins and senescence associated proteins identified in male *MafA^S64F/+^* mice. E) CM from WT MAFA or MAFA^S64F^ expressing EndoC-βH2 cells were collected, purified and added to EndoC-βH2 cells cultured for 72 hours. SASP genes upregulated in male *MafA^S64F/+^* mice were not identified in EndoC-βH2 cells. However, novel, species-specific secretory senescence-associated factors were identified in this context. N.D., not detectable. p*<0.05; **p<0.01. F) Schematic of temporal senescence and aging responses in *MafA^S64F^* expressing male β cells. In male *MafA^S64F/+^*, transiently high MafA protein triggers molecular insults to induce premature senescence marked by cell cycle arrest, senescence staining (blue cytoplasmic shade) and initiation of a senescence-associated secretory phenotype (SASP, pink secretory factors), resulting in impaired GSIS. Late (progressive) senescence propagates this phenotype with SASP amplification and diversification.

Progression of senescence includes development of SASP factors, which upon secretion induce senescence by reprogramming neighboring cells in a cell non-autonomous manner (Gorgoulis et al., 2019; Herranz and Gil, 2018). To determine if SASP factors are released by human cells expressing MAFA^S64F^, control medium or conditioned media (CM) collected from EndoC-βH2 cells expressing either WT MAFA or MAFA^S64F^ was applied to untransduced EndoC-βH2 cells (**Figure 7E**). Notably, SASP genes identified in male *MafA^S64F/+^* mice were not increased in EndoC-βH2 cells in response to WT MAFA or MAFA^S64F^ CM; however, human-specific SASP markers in related gene families to those identified in *MafA^S64F/+^* male mice were specifically increased by MAFA^S64F^ CM (**Figure 7E**). These results illustrate the ability of MAFA^S64F^ to promote cellular senescence in human β cells and to generate a functional, human β cell SASP via species-specific mediators.

## Discussion

Post-translational modifications of the key, islet-enriched MAFA protein are critical for regulating its activity, stability and cellular localization for appropriate β cell function (Guo et al., 2009; Guo et al., 2010; Han et al., 2007; Iacovazzo et al., 2018; Rocques et al., 2007). Here we have developed a mouse model of the human MAFA^S64F^ variant to understand mechanistically the sex-biased pathophysiological outcomes of MODY or insulinomatosis in affected heterozygous human carriers (Iacovazzo et al., 2018). Significantly, the physiological outcomes of *MafA^S64F/+^* mice appear to mimic the outcomes expected in human subjects, with glucose intolerance almost exclusively in males and improved glucose clearance and hypoglycemia in females.

This mutation blocks a key priming phosphorylation event at S65 which normally directs post-translational modifications impacting MAFA protein stability (Guo et al., 2009; Han et al., 2007; Rocques et al., 2007), transactivation (Han et al., 2007; Rocques et al., 2007), oncogenesis (Rocques et al., 2007), and DNA binding (Guo et al., 2010). These modifications are coupled to two antagonistic regulatory processes: increased transactivation activity and ubiquitin-mediated degradation (Rocques et al., 2007), illustrating that impeccable regulation of this protein is linked to islet β cell health. Earlier *in vitro* analysis demonstrated that the S64F MAFA mutation converted this normally unstable protein (t_1/2_ ~30 min) to a very stable form (t_1/2_-≥ 4 hours) (Iacovazzo et al., 2018). However, both male and female *MafA^S64F/+^* mice had similar improvements in glucose tolerance at 4 weeks, prior to overt and transient elevation in protein levels in just males (**Figures 1** and **2**). Notably, this increase in MafA^S64F^ protein levels preceded glucose intolerance seen by 5 weeks of age in males, while females continued to be modestly hypoglycemic with improved glucose clearance. The changes in male and female *MafA^S64F/+^* β cell activity were maintained throughout the period of analysis despite the presence of WT-like protein levels after 5 weeks. However, we propose that MafA^S64F^ is more abundantly and persistently produced than WT MAFA throughout the lifetime of the β cell despite similar protein levels, as supported by 3- to 5-fold lower *MafA* mRNA levels in male and female MafA^S64F/+^ islets in relation to the WT islets (**Figure 2C**). In line with this observation, human MAFA protein levels were largely unchanged upon comparing immunohistochemical staining in islets to the β cell mass in MAFA^S64F^ patients with insulinomatosis (Iacovazzo et al., 2018).

Our examination of ovariectomized female *MafA^S64F/+^* mice suggested that the estrogen sex hormone did not have a direct regulatory role, so it is presently unclear what factors are controlling the sex-biased phenotypes of MafA^S64F^ mutant mice. Since S64F MAFA-induced disease is not observed until around 38 years of age in both sexes without reports of affected pubertal development (Iacovazzo et al., 2018), we believe that neither estrogen nor testosterone are impactful to the divergent disease processes in humans. Instead, we propose that future efforts focus of determining whether sex chromosome gene expression is influential, such as the X chromosome-linked candidate genes found differentially expressed in 5-week-old *MafA^S64F/+^* male islets but not females compared to their respective WT controls (**Supplemental Figure 12**).

Bulk RNA-Seq analysis of 5-week-old male and female *MafA^S64F/+^* mouse islets illustrated similar dysregulation of genes involved in β cell identity and ion channel function (**Figure 3** and **Supplemental Figure 13**). For example, expression of several markers of β cell maturation were diminished in both *MafA^S64F/+^* male and female islets (**Supplemental Figure 13**), including the Mnx1 (Pan et al., 2015) and Pdx1 transcription factors as well as well the UCN3 neuropeptide (van der Meulen et al., 2015). In addition, expression of both sex-dependent and sex-independent genes involved in Ca^+2^ responses were identified (**Figure 3**). Accordingly, functional analyses after glucose stimulation identified dysfunctional Ca^+2^ responses. While all male 5-week-old mutant islets appear to have severely blunted glucose-induced Ca^+2^ responses (termed “non-responders” in **Figure 4**), females contained both “non-responder” and unique “responder” islets with higher frequency oscillations in response to glucose. Presumably, “responder” islets mediate the high basal glucose-induced insulin secretion properties characteristic of female *MafA^S64F/+^* islets (**Figure 4A** and **Supplemental Figure 3**), while downstream effectors of Mnx1 and Pdx1 activation that are involved in cell signaling (e.g. *Gcgr, Glp1r*), secretion (*Syt2, Ucn3, Ins1, Ins2*) and ion channel activity contribute to β cell non-responder dysfunction (Blum et al., 2014; Gilbert and Blum, 2018; Jacobson and Shyng, 2020; Kalwat and Cobb, 2017).

Because of the heterogeneity of the Ca^+2^ signaling changes in female *MafA^S64F/+^* islets, in this study we narrowed our focus to determine possible mechanisms contributing to the MafA^S64F^ induced diabetic phenotype in males. Future single cell sequencing efforts could reveal the factors regulating the interplay of distinct “responder” and “non-responder” islet β cell populations of *MafA^S64F/+^* females as well as their relative homogeneity in male non-responder islets. Importantly, our results clearly show that the many gene products associated with male *MafA^S64F/+^* β cell inactivity are not made in female variants (**Figure 5** and **Supplemental Figure 9**). As divergent regulation of metabolism between sexes in aging and disease is increasingly recognized (Sampathkumar et al., 2019), MAFA^S64F^ may provide a penetrant model to study sex-dependent effects on β cell health.

Since the compromised voltage-gated Ca^+2^ channel triggering of glucose-induced insulin secretion in *MafA^S64F/+^* males appeared to be caused by an inactivated β cell population, we focused on understanding how regulation was impacted in this context. Our results have demonstrated that MafA^S64F^ produces premature aging and senescence signatures in male, murine β cells and in the chromosomally male, human EndoC-βH2 cell line (**Figures 5, 6,** and **7**. In contrast, neither an aging or senescence signature was observed in female murine β cells expressing MafA^S64F^ (**Supplemental Figures 9** and **10**).

Unlike the temporary arrest of cellular quiescence, senescence is thought to be irreversible and refractory to mitogenic stimuli. Senescent cells undergo a progression of changes after early insult, including cellular arrest by activation of cell cycle inhibitors (i.e., p53, p21 and/or p16 (**Figures 5, 6** and **7**), among others) and DDR responses (Gorgoulis et al., 2019; Herranz and Gil, 2018). Progression to senescence involves chromatin remodeling to influence gene expression, metabolism, autophagy and SASP, with release of a heterogeneous mix of SASP effector proteins influencing neighboring cells in a cell non-autonomous manner (Gorgoulis et al., 2019; Herranz and Gil, 2018). Terminal senescence from persistent damage involves autocrine and paracrine SASP amplification, loss of nuclear integrity and diversification of the SASP phenotype (Gorgoulis et al., 2019). Notably, senescence-associated SA-β-gal staining was not induced in 4-week-old *MafA^S64F/+^* β cells, the time point just prior to detection of glucose intolerance (**Figures 1** and **6**).

Ultimately, senescent cells become resistant to apoptosis by up-regulation of anti-apoptotic proteins (such as those in the BCL2 family) and are often cleared by immune cells (Gorgoulis et al., 2019; Herranz and Gil, 2018). Interestingly, effective clearance of β cells is not apparent in *MafA^S64F/+^* male islets as we see accumulation of senescent cells, and there was no evidence of β cell death (**Figure 6**, **Supplemental Figure 11,** data not shown). Senescent cells can play a causal role in aging and aging-related pathology (van Deursen, 2014) and targeted removal of senescent cells can improve health span and reduce the incidence of aging-related diseases (Baker et al., 2016; Baker et al., 2011; Chang et al., 2016; Childs et al., 2016). Indeed, independent studies have shown that removal of the rare, senescent β cells in mouse models of T1D and T2D helps restore β cell function and glucose homeostasis *in vivo* (Aguayo-Mazzucato et al., 2019; Thompson et al., 2019). These senescent β cells showed distinct SASP signatures depending on modeling context (T1D vs. T2D) (Aguayo-Mazzucato et al., 2019; Thompson et al., 2019), and these signatures were also species-dependent such that only a subset of SASP mediators identified in respective mouse models were detected in human, diabetic islets (Aguayo-Mazzucato et al., 2019; Thompson et al., 2019).

Importantly, up to 50-60% of human T2D β cells were shown to be senescent (Aguayo-Mazzucato et al., 2019), compared to their rare occurrence in previously described mouse models (Aguayo-Mazzucato et al., 2019; Thompson et al., 2019). This rate of β cell senescence is comparable to those found in *MafA^S64F/+^* male islets, suggesting that this variant models the widespread β cell senescence in human T2D (**Figure 6**). In fact, MAFA^S64F^ expression induced β cell senescence in human EndoC-βH2 β cells with development of a functional SASP. Treatment with CM from MAFA^S64F^-expressing EndoC-βH2 cells induced β cell senescence and SASP factors different from those identified in *MafA^S64F/+^* male mouse islets, albeit from similar molecular families. In sum, these studies implicate MAFA^S64F^-induced senescence factors in causing β cell dysfunction and open doors to identify β cell senescence signatures across etiologies of diabetes (**Figure 7F**).

Human and rodent islets differ substantially in architecture, cell composition, proliferative capacity, islet amyloid formation, antioxidant enzyme levels, and, most significantly for the results described herein, MAFA and MAFB transcription factor expression (Bosco et al., 2010; Brissova et al., 2005; Butler et al., 2007; Cabrera et al., 2006; Dai et al., 2012; Fiaschi-Taesch et al., 2010; Henquin et al., 2006; Tyrberg et al., 2001). Because MafA is expressed at the onset of rodent β cell formation during embryogenesis but not until 10 years of age in humans (Cyphert et al., 2019; Dai et al., 2012; Hang and Stein, 2011), we appreciate that some of the observations made in *MafA^S64F/+^* mice will not be directly relevant to the human disease. For example, since substantial β cell proliferation ceases in humans prior to MAFA expression (Cyphert et al., 2019; Dai et al., 2012; Hang and Stein, 2011), the decreased islet cell proliferation observed in male and female *MafA^S64F/+^* mice will not be observed in humans. We also did not expect to find that *MafA^S64F/+^* male mice would manifest the overt fasting hyperglycemia seen in affected humans (Iacovazzo et al., 2018), since only one of the seven human MODY transcription factors has a phenotype in heterozygous mice comparable to human carriers (i.e., Pdx1 (Ahlgren et al., 1998)). As found in other studies of MODY transcription factor function in mice, only the homozygous (male) *MafA^S64F/S64F^* mice manifested an explicit elevation in fasting blood glucose levels and glucose intolerance, which worsened with age (**Supplemental Figure 2**). MafA^S64F^ presumably represents another example of species-specific differences in how gene dosage of critical regulatory gene variants impacts islet cell health.

It is also likely that the unique requirement of MAFB in human (but not rodent) β cells influences the effects of S64F MAFA in human carriers. The human MAFA^S64F^:MAFB heterodimeric activator could impart a unique influence on β cells compared to the mouse MafA^S64F^:MafA homodimeric activator (Cyphert et al., 2019; Hang and Stein, 2011), which may explain why neuroendocrine tumors and overt hypoglycemia was not observed in aged MafA^S64F/+^ mice (data not shown). Notably, MAFB was recently shown to be essential for the formation of human embryonic derived β cells and insulin production (Russell et al., 2020), whereas there is no phenotype associated with the loss of MafB in mouse islet β cells except during pregnancy (Cyphert et al., 2019). Thus, we believe that it will be important to extend the analysis of MAFA^S64F^ control to human islets both acutely *in vitro* and chronically after transplantation of human pseudoislets into immunocompromised mice to directly determine its effect *in vivo*. Such studies should generate keen insight into unique, species-specific, age-dependent and sex-biased molecular and genetic mechanisms controlling human islet β cell activity.

## Acknowledgements

This research was performed using resources and/or funding provided by the NIH-grants to R.S. (DK090570), E.M.W. (F32 DK109577), J.C. (T32 DK007061), F.M.J. (DK074970 and DK107444), J.S. (DK109102, HL144846), D.J. (DK097392), and the Vanderbilt Diabetes Research and Training Center (DK20593). X.T. was supported by a JDRF Fellowship (3-PDF-2019-738-A-N), J.C. by a Doris Duke Charitable Foundation Physician Scientist Fellowship and VUMC Harrison Society funds, D.I. by a George Alberti Research Training Fellowship funded by Diabetes UK (16/0005395), and F.M.J. also by a U.S. Department of Veterans Affairs Merit Review Award (BX003725). We also thank Drs. Raphaël Scharfmann and Phillippe Ravassard for generously providing EndoC-βH2 cells. Imaging was performed with NIH support from the Vanderbilt University Medical Center Cell Imaging Shared Resource (National Cancer Institute grant CA-68485; NIDDK grants DK20593, DK58404, and DK59637; Eunice Kennedy Shriver National Institute of Child Health and Human Development grant HD-15052; and National Eye Institute grant EY08126). Islet hormone analysis was performed in the Vanderbilt University Medical Center Islet Procurement and Analysis Core (NIDDK grant DK20593) and Vanderbilt University Neurochemistry Core (NICHD grant U54 HD083211). *MafA^S64F^* mice were produced in the Diabetes Research Center Animal Studies Core at the University of Michigan, which is supported by NIH grant P30 DK020572.

All authors have no conflict of interest to declare.

## Author Contributions

E.M.W., J.C., X.T. and R.S. designed the initial experiments. E.M.W., J.C., X.T. M.G. and J.-H.L. executed and analyzed the experiments with input from S.Y., D.I., S.F, M.K., S.K., F. M-J., D.J. and R.S. The manuscript was principally written by E.M.W., J.C., X.T. and R.S., although all authors have reviewed versions. R.S. is the guarantor of this work and, as such, had full access to all data in the study and takes responsibility for the integrity of the data and the accuracy of the data analysis.

