## Supplemental Figures for "Sex-biased islet β cell dysfunction is caused by the MODY MAFA S64F variant by inducing premature aging and senescence in males"

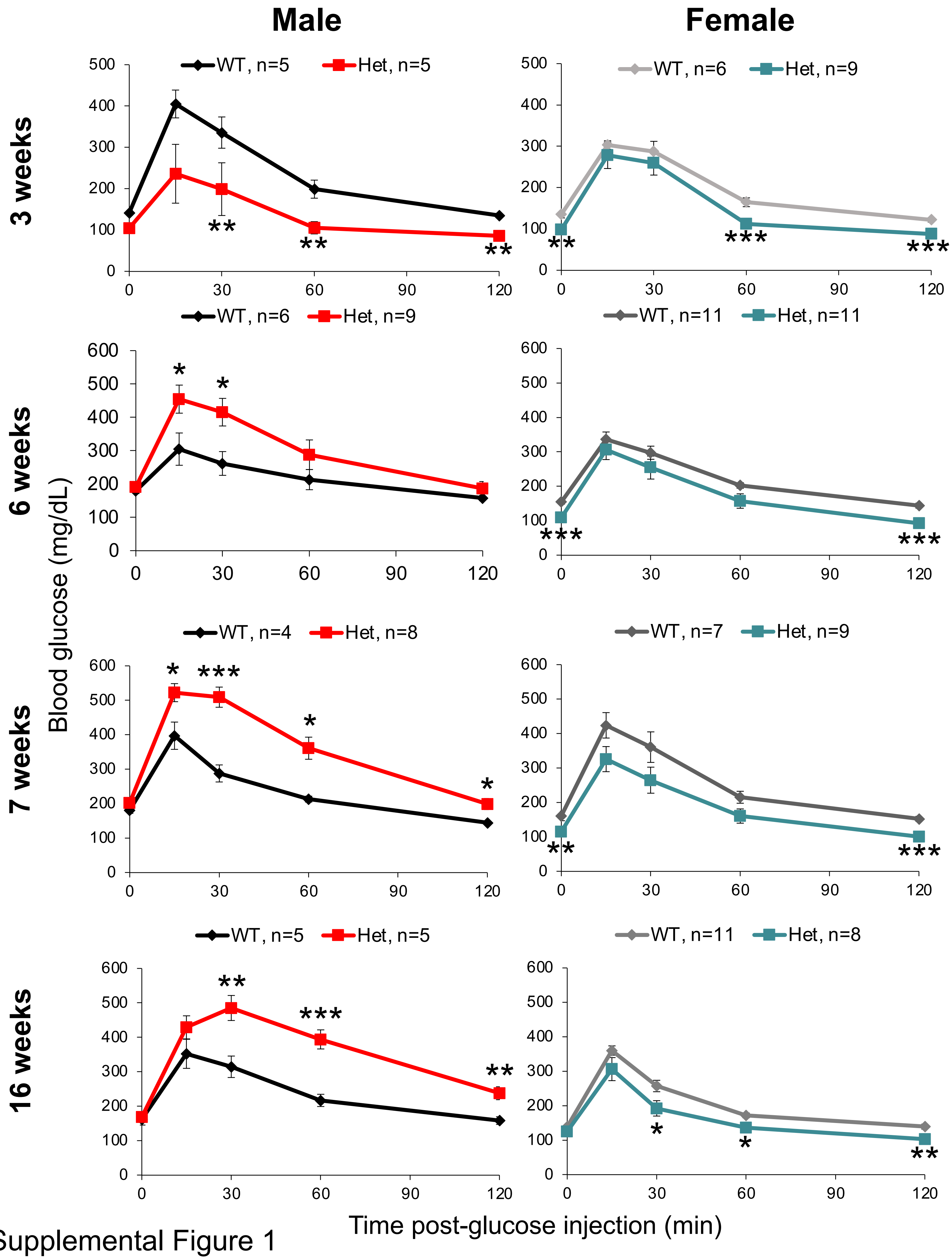

Supplemental Figure 1

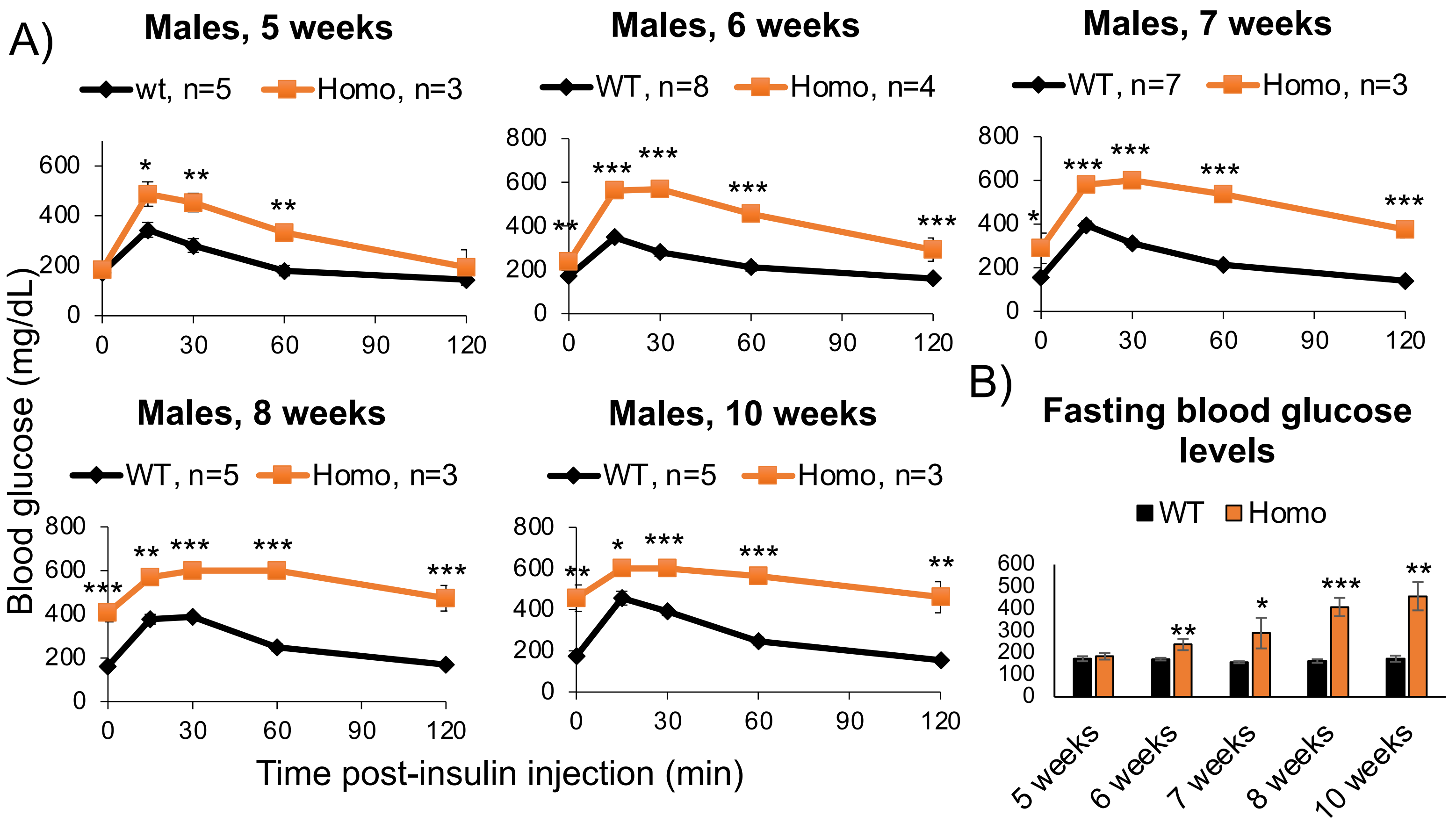

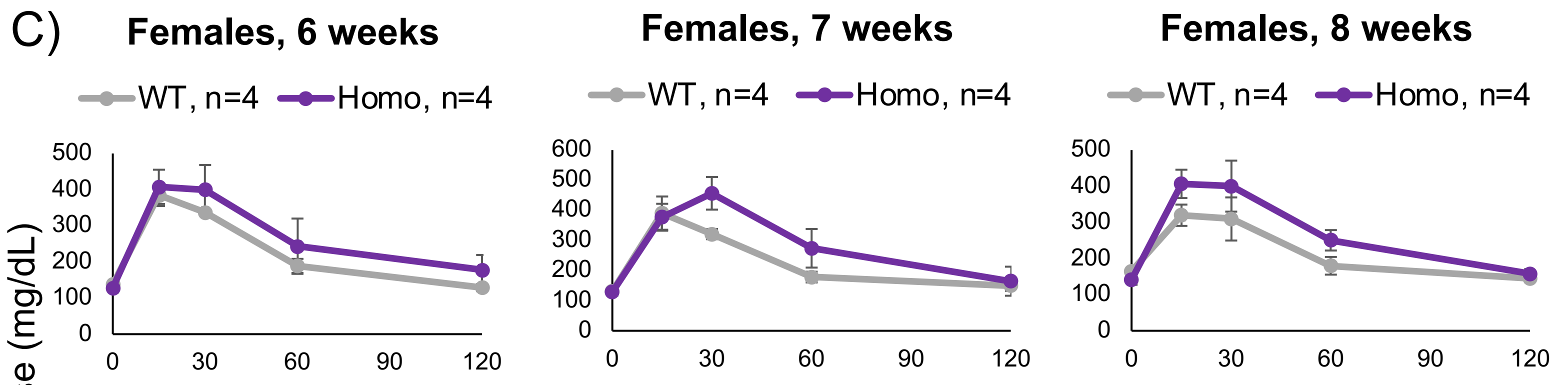

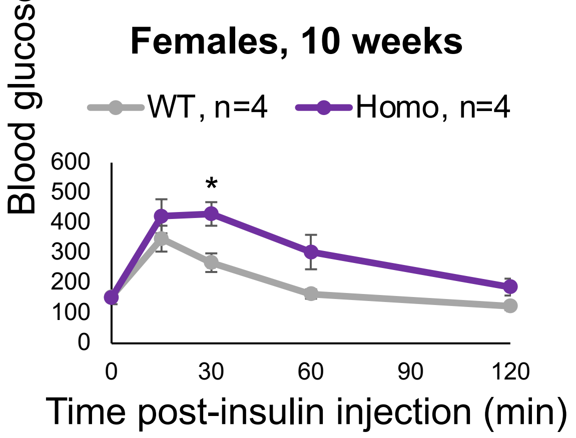

D)

| | Observed | Expected | $Chi^2$ p-value |
|---|---|---|---|
| WT | 57 | 37.5 | p<0.01 |
| Het | 82 | 75 | n.s. |
| Homo | 11 | 37.5 | p<0.001 |

Supplemental Figure 2

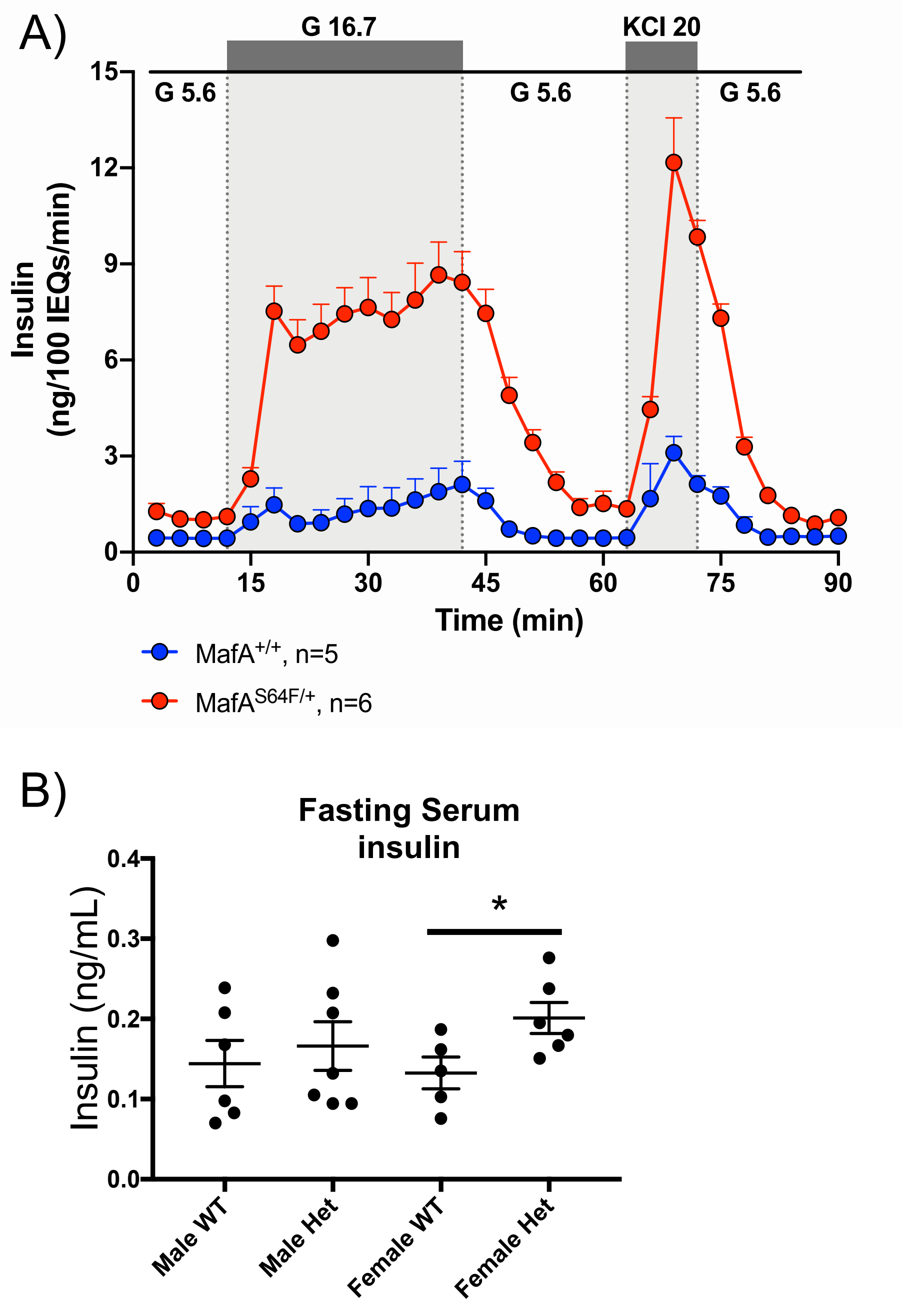

Supplemental Figure 3

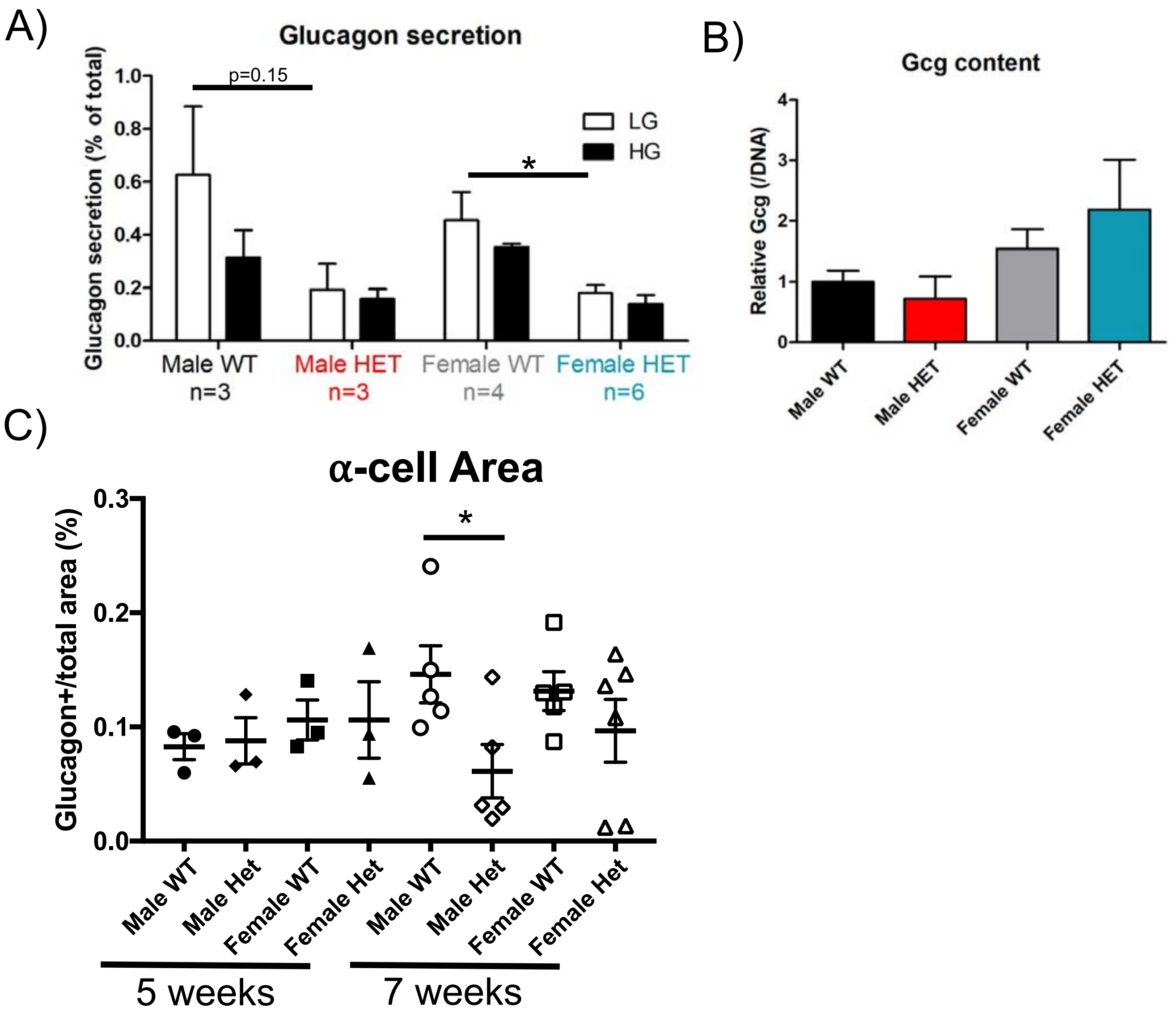

Supplemental Figure 4

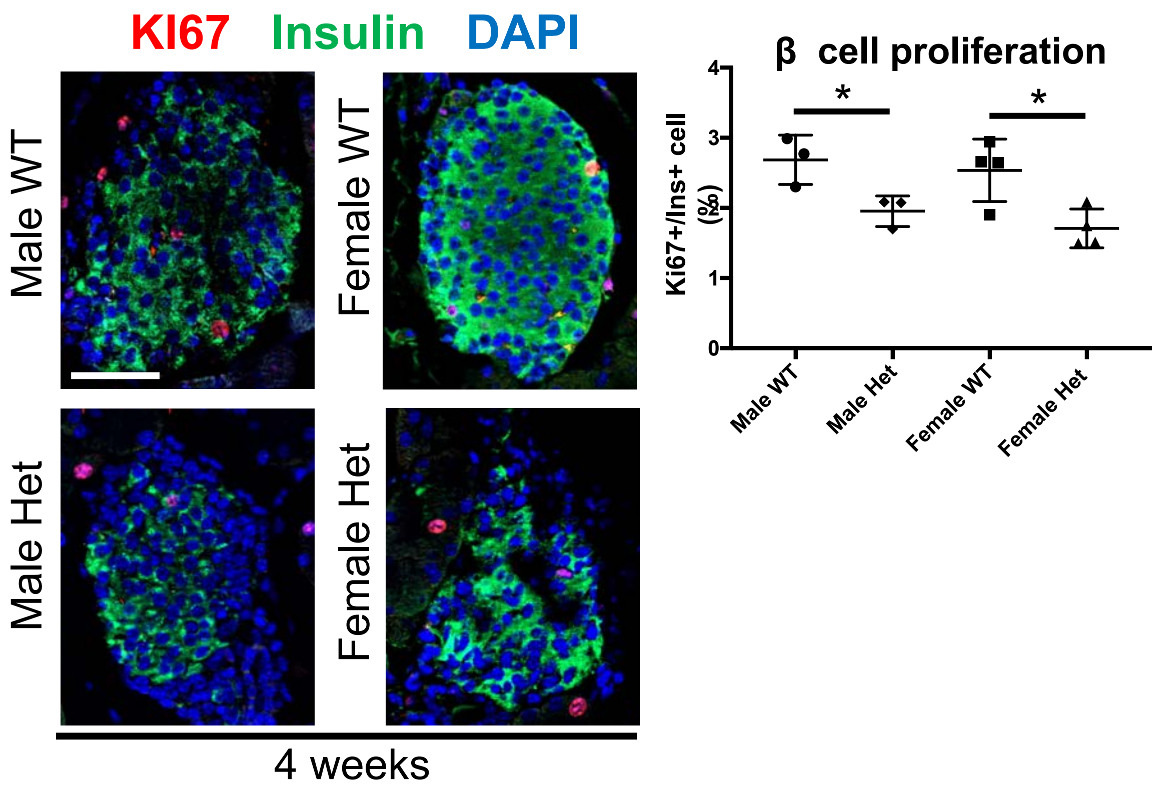

Supplemental Figure 5

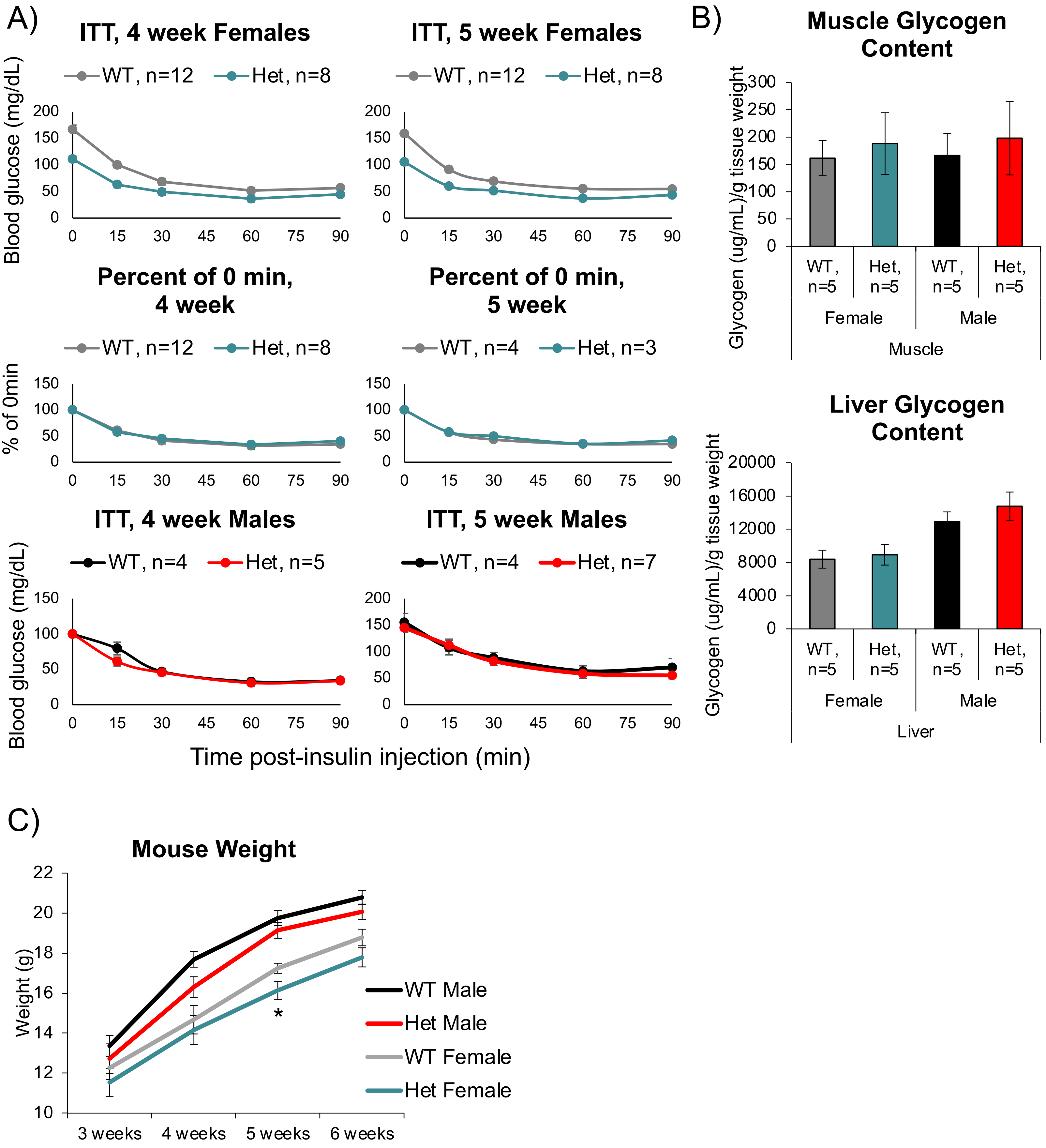

Supplemental Figure 6

A)

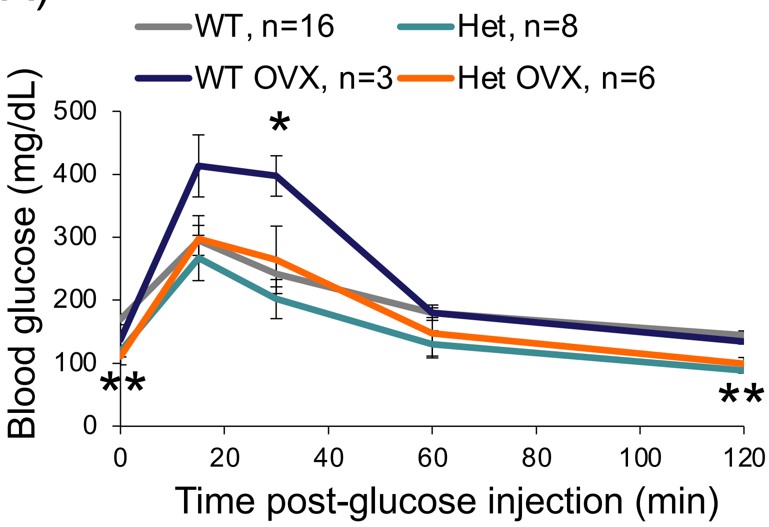

B)

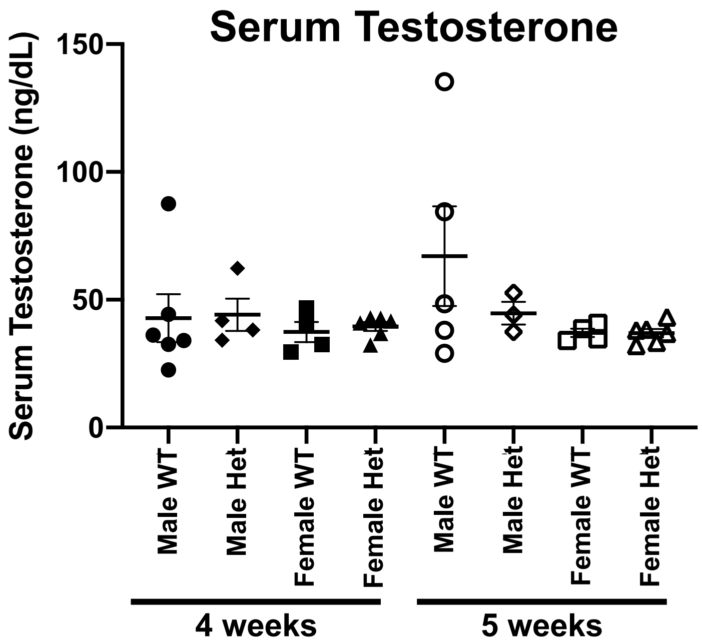

Supplemental Figure 7

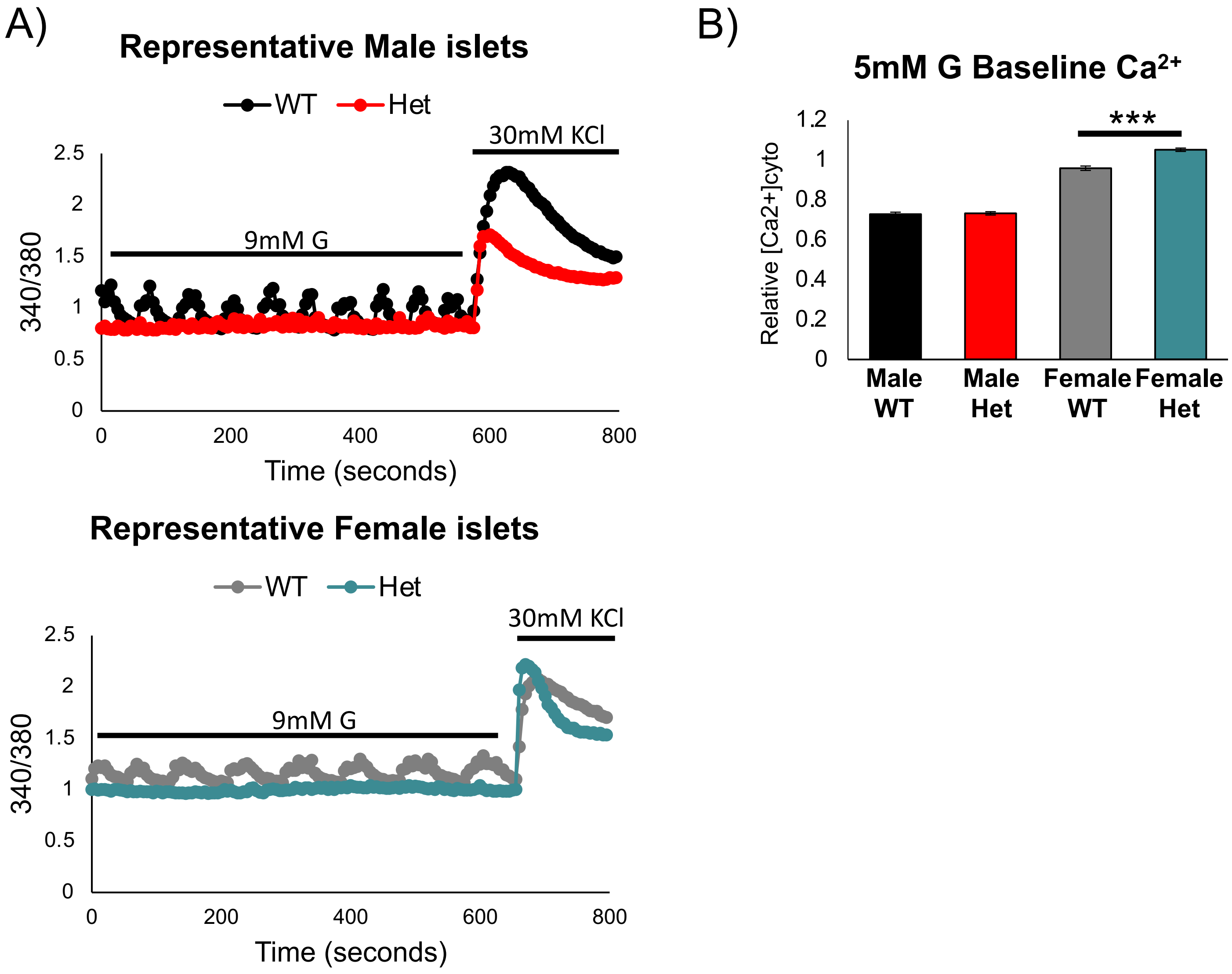

Supplemental Figure 8

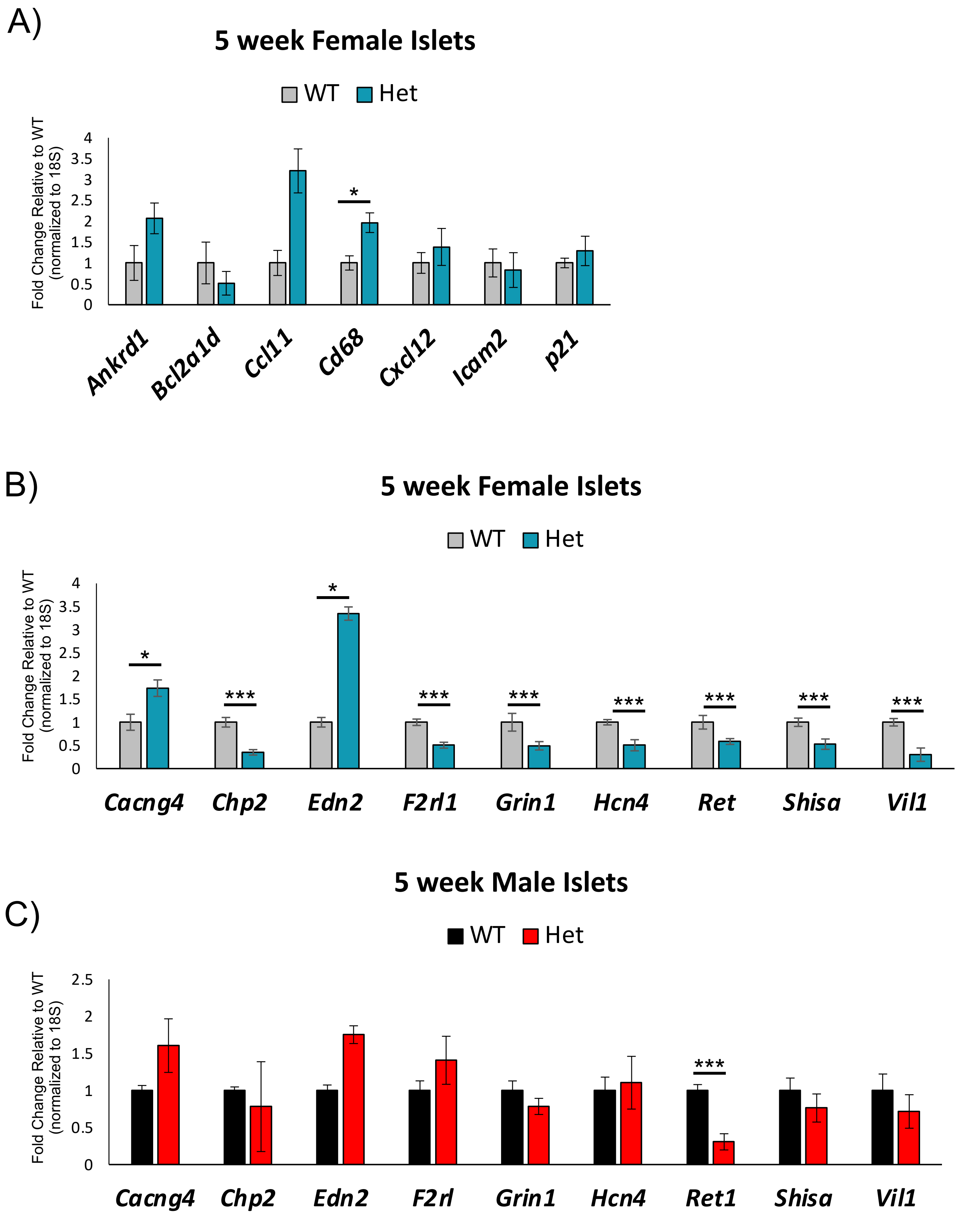

Supplemental Figure 9

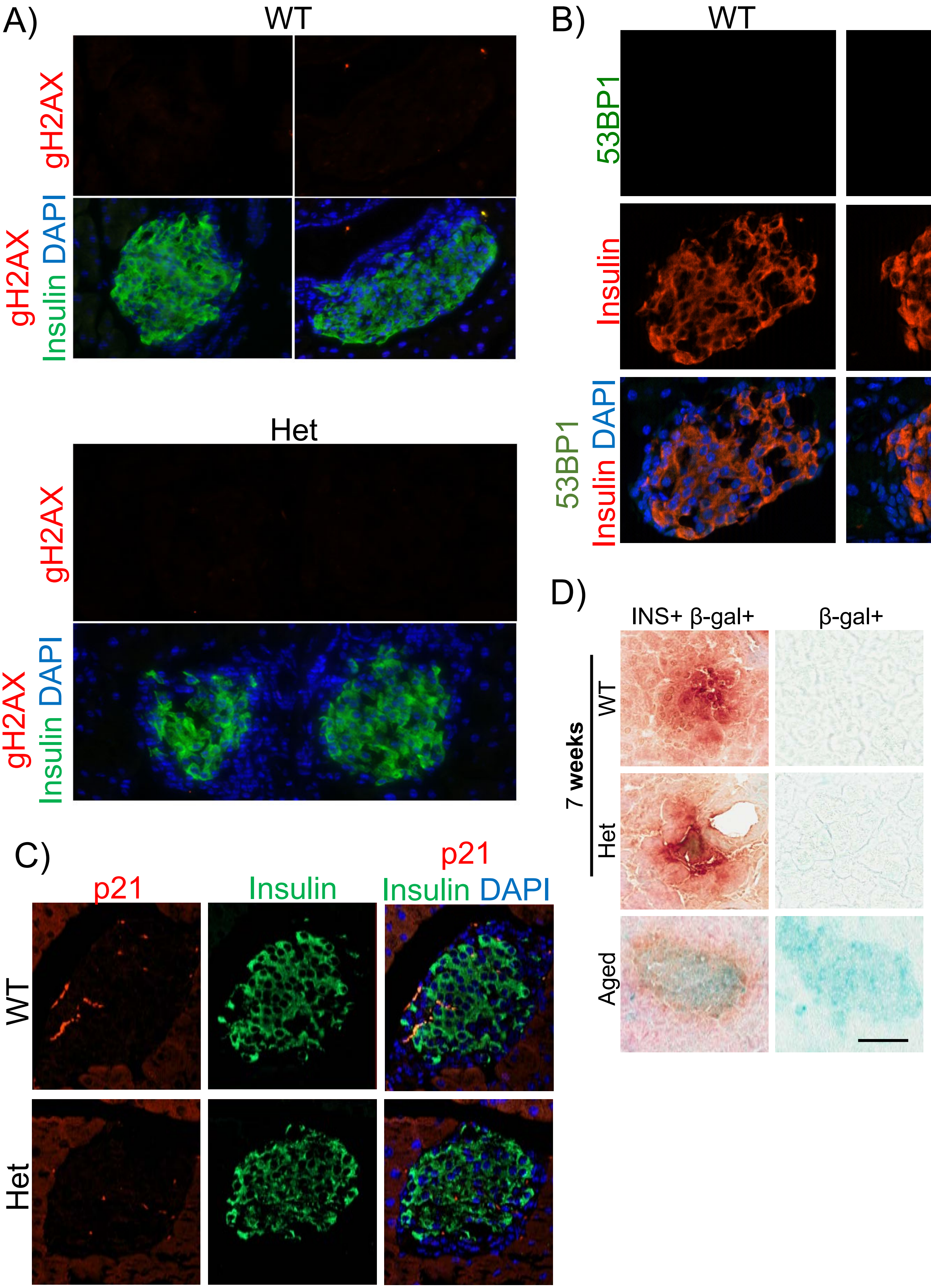

Supplemental Figure 10

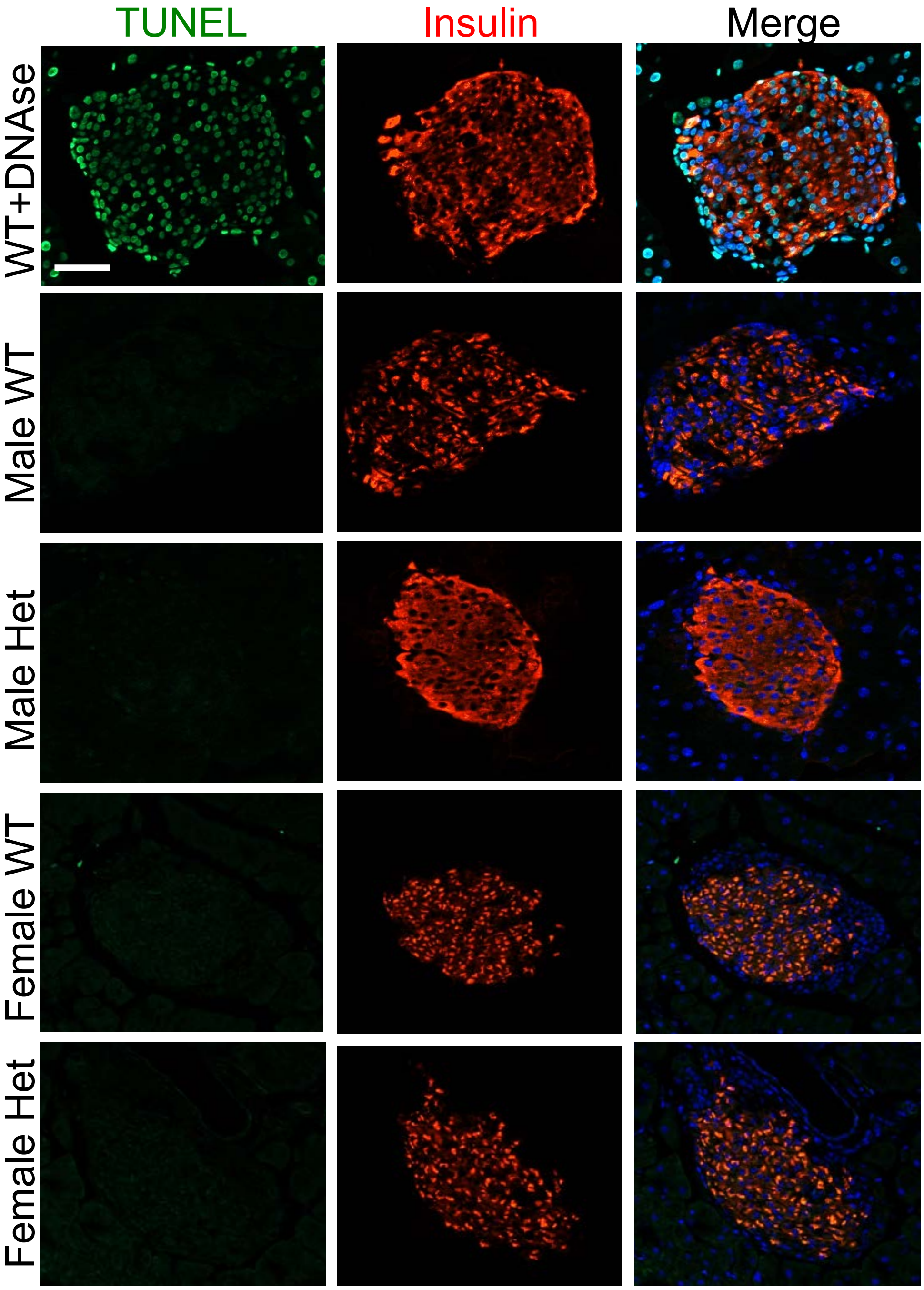

Supplemental Figure 11

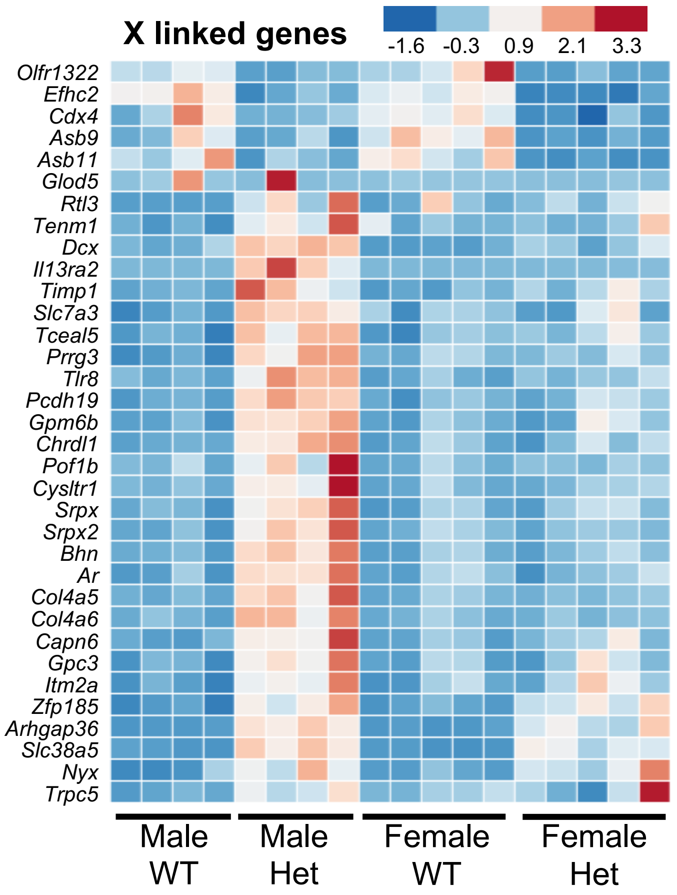

Supplemental Figure 12

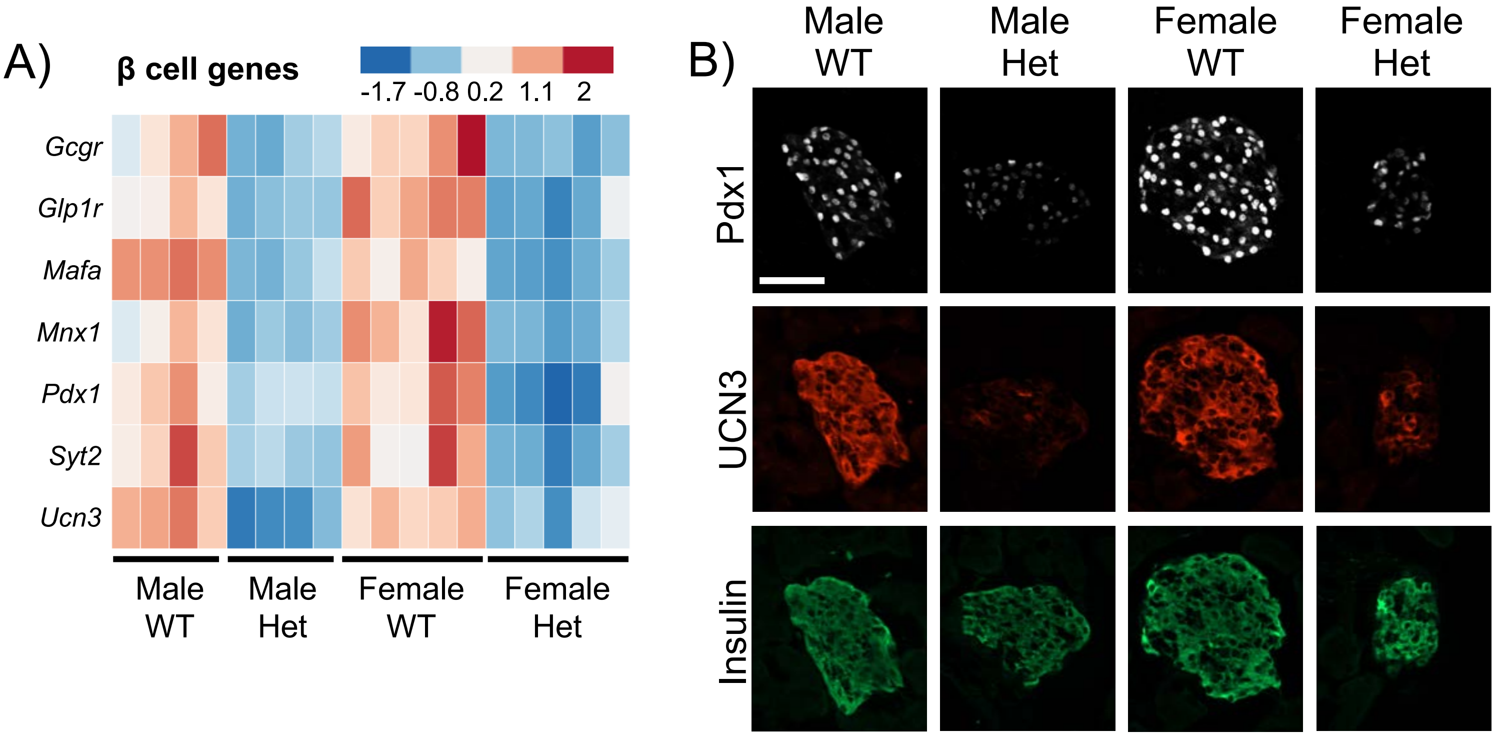

Supplemental Figure 13

Supp Table 1: Antibody information

| Antibody | Host species | Source | Catalog # | Application | Concentration |
|---|---|---|---|---|---|
| Insulin | guinea Pig | Thermo Fisher | PA1 26938 | IF/IHC | 1 to 500 |
| Glucagon | mouse | Sigma | G2654 | IF/IHC | 1:4000 |
| Ki67 | mouse | BD Biosciences | 550609 | IF | 1:1000 |
| DAPI | n/a | SouthernBiotech | 0100-20 | IF | n/a |
| MAFA | rabbit | Cell Signaling | 79737 | IF | 1:20,000 |
| Pdx-1 | goat | Wright lab | n/a | IF | 1:20,000 |
| UCN3 | rabbit | Phoenix | H-019-29 | IF | 1:500 |
| p21 | rat | abcam | ab107099 | IF | 1:200 |
| 53BP1 | rabbit | Bethyl | A300-272A | IF | 1:400 |
| γH2AX | rabbit | abcam | ab81299 | IF | 1:4000 |
| Tunel assay kit: "In Situ Cell Death Detection Kit Fluorescein", (Roche, Cat# 11684795910) | | | | | |

### Supp Table 2: Primer information

| Mouse primers | Forward | Reverse |
|---|---|---|
| *MafA* | CCTGTAGAGGAAGCCGAGGAA | CCTCCCCCAGTCGAGTATAGC |
| *Ins1* | CACTTCCT ACCCCTGCTGG | ACCACAAAGATGCTGTTTGACA |
| *pre-Ins2* | GGGGAGCGTGGCTTCTTCTA | GGGGACAGAATTCAGTGGCA |
| *Ins2* | CCACCCAGGCTTTTGTCAAA | CCCAGCTCCAGTTGTTCCAC |
| *Pdx1* | CGGCTGAGCAAGCTAAGGTT | TGGAAGAAGCGCTCTCTTTGA |
| *Ucn3* | AGCACCCGGTACAGATACCAA | GGCCTTGTCGATGTTGAAGAG |
| *Ankrd1* | AAACGGACGGCACTCCACCG | CGCTGTGCTGAGAAGCTTGTCTCT |
| *Bcl2a1d* | GTATATCCACTCCCTGGCTGAG | TAGTCACAATCCTTCCCCAGTT |
| *Ccl11* | TCCACAGCGCTTCTATTCCTG | GGAGCCTGGGTGAGCCA |
| *Cd68* | ACTTCGGGCCATGTTTCTCT | GCTGGTAGGTTGATTGTCGT |
| *Cxcl12* | CAGTGACGGTAAACCAGTCAGC | TGGCGATGTGGCTCTCG |
| *Icam1* | CAATTTCTCATGCCGCACAG | AGCTGGAAGATCGAAAGTCCG |
| *p21* | GCCTTAGCCCTCACTCT TG | AGCTGGCCTTAGAGGTGACA |
| *Cacng4* | TCCGGAAGACGCGGACTAC | ATGATGTTGTGGCGTGTCTTG |
| *Chp2* | CGCCTAGACCTCCAGCAGATC | GCCTGCGAAATACAGTCTCTGAC |
| *Edn2* | CTGGCAAGATGTGGACTGCTGA | GCCTTTCTTGTCACCTCTGGCT |
| *F2rl1* | CGGACCGAGAACCTTGCACCG | GTGAGGATGGACGCAGAGAACT |
| *Grin1* | CCTTTCAGAGCACACTGTGGCT | CCAGGAAAACCACATGGCAGAG |
| *Hcn4* | CGTGCTCACTAAGGGCAACAAAG | GCACCTCATTGAAGTTGTCCACG |
| *Ret* | TTCCAGCATCAACTGCACTG | GTCAGTGGCTACCACCGTGT |
| *Shisa2* | TGCGACAACGACCGCCAGCAG | TGAAGGCAACGAACACTGAGCC |
| *Vil1* | TTCTACGGTGGTGACTGCTACC | TGGTCCAACAGGACGGCTTGAT |
| *β-Actin* | AGGTCATCACTATTGGCAACGA | CACTTCATGATGGAATTGAATGTAGTT |

| Human primers | Forward | Reverse |
|---|---|---|
| *MAFA* | GAGAGCGAGAAGTGCCAACT | TTCTCCTTGTACAGGTCCCG |
| *MAFB* | CATAGAGAACGTGGCAGCAA | ATGCCCGGAACTTTTTCTTT |
| *INS* | AGAGGCCATCAAGCAGATCACTGT | ACAGGTGTTGGTTCACAAAGGCTG |
| *ANKRD1* | AGACTCCTTCAGCCAACATGATG | CTCTCCATCTCTGAAATCCTCAGG |
| *BCL2A1* | GGATAAGGCAAAACGGAGGCTG | CAGTATTGCTTCAGGAGAGATAGC |
| *CCL11* | GCTACAGGAGAATCACCAGTGG | GGAATCCTGCACCCACTTCTTC |
| *CD68* | CGAGCATCATTCTTTCACCAGCT | ATGAGAGGCAGCAAGATGGACC |
| *CXCL12* | CTCAACACTCCAAACTGTGCCC | CTCCAGGTACTCCTGAATCCAC |
| *ICAM1* | AGCGGCTGACGTGTGCAGTAAT | TCTGAGACCTCTGGCTTCGTCA |
| *P21* | CCT GTC ACT GTC TTG TAC CCT | GCG TTT GGA GTG GTA GAA ATC T |
| *BCL2* | ATCGCCCTGTGGATGACTGAGT | GCCAGGAGAAATCAAACAGAGGC |
| *CXCL2* | GGCAGAAAGCTTGTCTCAACCC | CTCCTTCAGGAACAGCCACCAA |
| *ICAM3* | AGATCGTCTGCAACGTGACCCT | TCGCTGAGGTTCACAATGGGTC |
| *IGFBP2* | CGAGGGCACTTGTGAGAAGCG | TGTTCATGGTGCTGTCCACGTG |
| *IGFBP4* | GAGCTGGGTGACACTGCTTG | CCCACGAGGACCTCTACATCA |
| *IGF1R* | CTCCTGTTTCTCTCCGCCG | ATAGTCGTTGCGGATGTCGAT |
| *GAPDH* | CTCACCGGATGCACCAATGTT | CGCGTTGCTCACAATGTTCAT |

**Supplemental Figure 1: Glucose tolerance phenotypes were stably maintained in male and female *MafA$^{S64F/+}$* mice.** GTT was performed in 3-, 6-, 7-, and 16-week-old mice. Glucose (2mg/kg) was injected following a 6 hour fast and blood glucose was measured at the indicated time points. *p<0.05; **p<0.01; ***p<0.001

**Supplemental Figure 2: Homozygous male *MafA$^{S64F/S64F}$* mice were hyperglycemic, with glucose tolerance and blood glucose levels worsening with age.** A) *MafA$^{S64F/S64F}$* mice showed progressively worsening glucose tolerance and fasting blood glucose levels between 5 to 10 weeks of age. B) Fasting blood glucose levels in homozygous male mutant mice increased significantly over time. C) Female S64F MafA homozygous mice were only mildly glucose intolerant at 10 weeks of age although their temporal responses to glucose was variable between animals. D) Chi-square analysis revealed that *MafA$^{S64F/S64F}$* male and female animals were observed with significantly less frequency at weaning than WT animals. In contrast, *MafA$^{S64F/+}$* numbers were as expected. *p<0.05; **p<0.01; ***p<0.001.

**Supplemental Figure 2: Female *MafA$^{S64F/+}$* islets secreted more insulin in response to low or high glucose.** A) Islet perifusion demonstrated islets from *MafA$^{S64F/+}$* female mice secreted higher levels of insulin at low (G 5.6) and high (G 16.7) glucose and in response to KCl. B) Serum insulin levels (ng/mL) were increased in 6-hour fasted female Het animals while male Het levels were unchanged.

**Supplemental Figure 4: Glucagon secretion was compromised in male and female *MafA$^{S64F/+}$* mice.** A) Glucagon secretion was decreased in S64F MafA Het islets, which were were incubated in 4.6 mM (LG) and 16.7 mM (HG) glucose for 1 hour prior to collection. Secretion was normalized to glucagon content (B), which was unchanged. Content was normalized to DNA. C) Islet $\alpha$ cell area was reduced in male S64F MafA Het mice, though measurements were highly variable between samples. Islet $\alpha$ cell area was calculated by dividing the total glucagon$^{+}$ area by the total pancreas area (eosin staining) multiplied by 100 to obtain percent (%). *p<0.05.

**Supplemental Figure 5: Islet β cell proliferation at 4 weeks of age was reduced in both male and female *MafA$^{S64F/+}$* mice.** A) Representative islets stained for insulin, Ki67 (proliferation marker), and DAPI (nuclei). Islet β cell proliferation was calculated by dividing the number of Ki67$^{+}$ cells by total insulin$^{+}$ β cells. Greater than 1000 β cells were counted per sample. *p<0.05.

**Supplemental Figure 6: Insulin tolerance and glycogen content are unaltered in *MafA$^{S64F/+}$* animals.** A) Insulin tolerance was unchanged at 4 and 5 weeks in Het animals once corrected for initial fasted blood glucose levels (Percent of 0 min). Insulin (0.5U/kg body weight) was injected following a 6 hour fast. B) Glycogen content in muscle (soleus) and liver was also unaffected in 7 week-old S64F MafA Het mice. Glycogen content was normalized to wet tissue weight. C) There was no change in average body weight between 3-6 weeks except a small decrease (-1.07 fold) in female S64F MafA Het mice at 5 weeks of age. N=5/group, *p<0.05.

**Supplemental Figure 7: Ovariectomy (OVX) did not alter glucose tolerance in female *MafA$^{S64F/+}$* mice, nor are testosterone levels changed.** A) Compared to 4 week-old control mice (i.e. non OVX: WT, grey line; Het, teal line), the GTT at 1-week post ovariectomy showed glucose intolerance in WT female mice (WT OVX, purple line) and no change in S64F MafA Het. *p<0.05; **p<0.01. B) Serum testosterone levels were unchanged in male or female S64F MafA Het. Male and female testosterone levels are similar at 4 and 5 weeks of age.

**Supplemental Figure 8: Male *MafA$^{S64F/+}$* islets had a reduced response to KCl stimulation, while female Het islets had an increased KCl response and baseline $Ca^{2+}$.** A) Representative $Ca^{2+}$ traces quantitated in Figure 4C in response to both high glucose (9 mM) and KCl (30 mM). B) Female S64F MafA Het islets have increased baseline $Ca^{+2}$ at 5mM compared with WT islets (right) while male Het islets do not (left).

**Supplemental Figure 9: qPCR confirmation of changes in islet gene expression identified by RNA-seq.** A) Senescence markers were primarily unchanged in female *MafA*$^{S64F/+}$ islets. B) Gene altered specifically in female *MafA*$^{S64F/+}$ islets identified by RNA-seq were unchanged in male *MafA*$^{S64F/+}$ islets (C). *p<0.05; ***p<0.001.

**Supplemental Figure 10: DNA damage and cell cycle inhibition markers were not detected in 5-week-old female *MafA*$^{S64F/+}$ islets.** Immunostaining for γH2AX and 53BP1 (DNA double strand break; A-B), p21 (cell cycle inhibitor; C), and endogenous SA-β-gal (senescence marker; D) were not detected in female Het islets.

**Supplemental Figure 11: Significant levels of apoptosis was not found in *MafA*$^{S64F/+}$ islet cells.** TUNEL$^{+}$ nuclei were barely detected in S64F MafA Het islets at 6 weeks of age but easily visible in DNAse treated controls (top panels).

**Supplemental Figure 12: Many X chromosome linked genes were altered in male *MafA*$^{S64F/+}$ islets.** Heat map developed from the 5-week-old RNA-Seq data. FDR<0.05.

**Supplemental Figure 13: Production of some β cell identity gene products are downregulated in *MafA*$^{S64F/+}$ islets.** A) Heat map showing β identity genes decreased in both male and female Het islets. FDR<0.05. B) Immunostaining illustrating decreased Pdx1 and UCN3 protein levels in 7-week-old S64F MafA Het islets.
